# Genetic coupling of hydathode formation with leaf morphogenesis maintains water homeostasis

**DOI:** 10.64898/2026.07.23.740248

**Authors:** Pauline Savourat, Anne-Sophie Sarthou, Panagiotis Papadopoulos, Damien Vasselon, Aurore Olympio, Néro Borrega, Stéphanie Pateyron, Christine Paysant Le Roux, Cléa Quentin, Jean-Marc Routaboul, Bernard Genty, Laurent Noel, Nathalie Leonhardt, Patrick Laufs

## Abstract

Hydathodes are specialized leaf structures present across vascular plants that allow guttation by connecting the xylem to the external environment through epithem tissue and permanently open water pores. However, the genetic mechanisms controlling their formation and physiological roles remain poorly understood. Here, we identify a genetic regulatory network that controls hydathode formation and links this process to leaf morphogenesis. This network converges on auxin signaling to coordinate the formation of the three hydathode cell types. We further show that epithem development requires sustained cell proliferation with limited endoreduplication. Analysis of hydathode mutants demonstrates that hydathode size and number are required to prevent reversible leaf flooding. Together, these findings establish a genetic framework for hydathode morphogenesis and uncover a central role for hydathodes in maintaining leaf water homeostasis under fluctuating environmental conditions.

## Introduction

Plants possess a remarkable diversity of secretory structures, including nectaries, salt or oil glands, and glandular trichomes, which release a wide range of compounds ^1–3^. Among these, hydathodes occur across all major groups of vascular plants ^4^ and display a tripartite organization consisting of (i) a group of small cells, the epithem, laying within the larger spongy mesophyll layer, (ii) xylem elements connecting the epithem to the vascular system, and (iii) permanently open water pores (or hydathode pores) that resemble enlarged stomata (Extended Data Fig. 1A-U, 2) ^5–7^. Hydathodes, which are often located at the apex of leaves or at the tip of serrations, like in *Arabidopsis thaliana* (Arabidopsis), establish a direct hydraulic connection between the plant vascular system and the external environment through the epithem and the water pores, enabling guttation, the exudation of xylem-derived fluids as small droplets (Extended Data Fig. 1V). Guttation occurs when the influx of xylem sap into the leaf exceeds its capacity to dissipate water through evapotranspiration. This situation typically takes place when elevated xylem root pressure is combined with reduced evapotranspiration resulting from stomata closure or high atmospheric humidity ^8,9^. To minimize the loss of valuable metabolites in the guttation fluid, hydathodes express a wide range of transporters that facilitate metabolite scavenging ^5,10^. However, because water pores remain permanently open, hydathodes also constitute privileged entry points for pathogens, and consequently, immune responses are activated at hydathodes to defend against microbial invasion ^7,11–13^. The physiological and ecological roles of hydathodes and guttation remain mostly elusive. Mechanical sealing of *Chloranthus japonicus* hydathodes prevents guttation and induces leaf flooding, the filling of intercellular airspaces by fluids, which severely impairs photosynthesis^14^. Additional functions have been proposed, including roles in pathogen dissemination or resistance, insect nutrition, redistribution of water from deep soil layers to the surface, or foliar water uptake ^15,16^.

Further progress in elucidating the roles of hydathodes is limited by the scarcity of well-characterized mutants defective for hydathode development. Indeed, the molecular, genetic, and, to a lesser extent, cellular processes governing hydathode formation remain poorly understood. In Arabidopsis, water pore formation relies on genes essential for proper leaf blade expansion ^17^. High auxin biosynthesis and signaling, mark the site of epithem formation from early developmental stages through to mature hydathodes ^10,18–20^. However, since this auxin distribution pattern is also required for the development of the tooth structure within which the hydathode forms ^21,22^, it remains unclear whether auxin accumulation actively and directly promotes hydathode formation or merely establishes a developmental context—the tooth—necessary for hydathode formation. The phenotype of the *yuc1246* quadruple mutant, defective in the auxin-biosynthetic YUCCA (YUC) enzymes, exemplifies this ambiguity: these plants form small, narrow leaves lacking teeth and exhibit abnormal hydathode development^20^. Consequently, the regulatory mechanisms specifically controlling hydathode formation remain unresolved. Here, we show that hydathode formation can be genetically uncoupled from tooth development in Arabidopsis and define a dedicated molecular framework that sequentially controls hydathode specification and differentiation. The identification of hydathode-defective mutants with divergent leaf morphologies genetically establishes hydathodes as critical components of plant water homeostasis.

## Results

### Hydathodes can be uncoupled from tooth fate

In serrated leaves such as Arabidopsis, hydathodes have been described as forming at the leaf apex and tooth tips ^23,24^. However, the number of marginal hydathodes was not strictly correlated to the number of leaf teeth (Extended Data Fig. 3A), and closer examination showed that hydathodes could also develop between two tooth tips along the margin of large, late forming rosette leaves. These structures showed the typical hydathode organization (connected to the vascular system by xylem cells, containing small epithem cells and covered by large water pores) and expressed several hydathode reporters, including E325-GFP (Fig. 1A-G, Extended Data Fig. 3B-D). To further question the relationship between hydathode and tooth fate, we investigated hydathode formation in *cup-shaped cotyledon 2* (*cuc2*) mutants that form smooth leaf margins with no developed teeth ^25^. In *cuc2-1* leaves, the formation of the apical hydathode was unaffected, whereas the number of marginal hydathodes was severely reduced. Nevertheless, some marginal hydathodes with normal cellular features, typical reporter gene expression, and guttation capacity still developed in *cuc2-1* leaves (Fig. 1H, J, M, Extended Data Fig. 3E, G-K). Because *CUC2* genetically interacts and plays redundant roles with the *DORNRÖSCHEN (DRN)* and *DORNRÖSCHEN-LIKE* (*DRNL*) genes during embryo and lateral organ development ^26–28^, we next investigated hydathode formation in *drn drnl* double mutants. The *drn-1 drnl-1* double mutant that formed leaves with fewer and less developed teeth, also showed a reduced number of hydathodes (Fig. 1I, K, L, Extended Data Fig. 3F). In some cases, a tooth without any clear hydathode formed on *drn-1 drnl-1* Ieaves (Fig. 1I). Because the *drn-1 drnl-1* double mutant is in a mixed Col/L*er* genetic background and could show variable leaf shape, we generated 3 novel *drnl-cr* alleles via genome editing in the Col-0 background (Extended Data Fig. 4A-D). The apical hydathode was absent in about 40% of the three new *drnl-cr* mutants and *drnl-cr4 drn-3* double mutants showed a very strong reduction in both apical and marginal hydathodes (Extended Data Fig. 4E, F). Hydathode number was not significantly modified in the single *drn-2* and *drn-3* mutants (Extended Data Fig. 4E).

**Figure 1.**
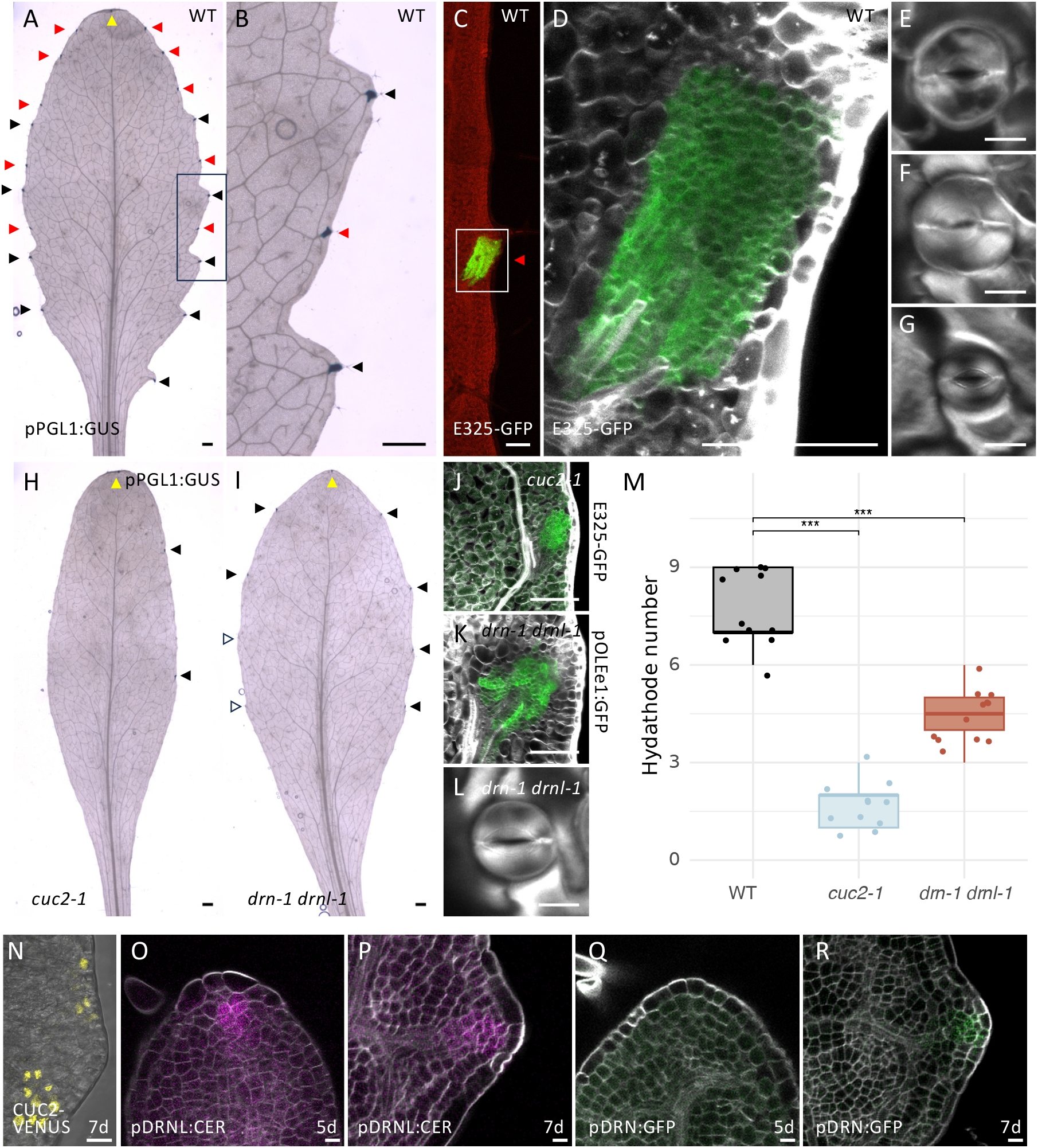
Hydathodes can be uncoupled from tooth fate. **A, B.** pPGL1:GFP-GUS expression in a wild-type leaf 11. **C, D.** E325:GFP expression in a hydathode located between two teeth in a wild-type leaf 11 **E**. Water pores observed on a wild-type apical hydathode **F.** Water pores observed on a wild-type hydathode located between two teeth **G**. Stomata observed on a wild-type leaf blade **H, I.** pPGL1:GFP-GUS expression in a *cuc2-1* and in *drn-1 drnl-1* leaf 11 **J.** E325-GFP expression in a marginal hydathode of a *cuc2-1* leaf **K.** pOLEe1:GFP-GUS expression in a marginal hydathode of a *drn-1 drnl-1* leaf **L.** Water pores observed on a marginal hydathode of a *drn-1 drnl-1* leaf **M.** Number of hydathodes in leaf 6 of WT, *cuc2-1* mutant and *drn-1 drnl-1* double mutant (*n* ≥ 11). **N.** pCUC2:CUC2:VENUS expression in developing tooth of leaf 1 at 5 days **O, P.** pDRNL:CER expression in the apex (**O**) or the developing tooth (**P**) of leaf 1 at 5 days (**O**) or 8 days (**P**) **Q, R.** pDRN:GFP expression in the apex (**Q**) or the developing tooth (**R**) of leaf 1 at 5 days (**Q**) or 7 days (**R**) Yellow arrow heads point to apical hydathodes, black arrow heads point to marginal hydathodes associated with a tooth while red arrow heads point hydathodes not associated with a tooth in **A**-**C**, **H** and **I**. GFP signal is shown in green in **C**, **D**, **J**, **K**, **Q** and **R,** CER signal is shown in magenta in **O** and **P**, VENUS signal is shown in yellow in **N**, fluorescence of the chlorophyl is shown in red in **C**, and grey shows SR2020 cell wall staining in **D-G, J**-**L, O-R** and shows transmitted light in **N**. Bars= 1 mm in **A**, **B**, **H** and **I**; 100 µm in **C**, **D**, **J** and **K**, and 10 µm in **E**-**G**, **K**, **L**, and **N-R**.

From these observations we first concluded that hydathodes formation can be genetically uncoupled from tooth fate in Arabidopsis. Second, the reduced hydathode number in *cuc2* and *drn dnrl* double mutants and the presence of teeth lacking hydathodes also showed that these genes contribute to hydathode formation. To further support this, we examined the expression patterns of these genes during hydathode development, using the first pair of leaves as a model system. In these leaves, the apical hydathode develops 2–3 days earlier than the two (or sometimes only one) marginal hydathodes (Extended Data Fig. 5). The pCUC2:CUC2:VENUS reporter was expressed around the developing tooth and incipient marginal hydathode site and no expression was observed at the leaf apex (Fig. 1N) ^21,22,29^. The pDRNL:CER reporter was transiently expressed at the incipient apical and marginal hydathode sites before any visible sign of epithem differentiation (Fig. 1O, P, Extended Data Fig. 6A, B), while the pDRN:GFP reporter was also expressed at the incipient marginal hydathode site, but was not detected at the leaf apex (Fig. 1Q, R, Extended Data Fig. 6C). The expression patterns of *DRN* and *DRNL* corroborate with the specific defect of apical hydathode formation observed in single *drnl* mutants and the defects in marginal hydathodes in *drn drnl* double mutants. The presence of an hydathode at the leaf apex of *cuc2* and *drn* mutants suggests that additional regulators may be acting at the leaf apex to promote hydathode formation.

### Transcriptomic profiling reveals two phases during hydathode formation

To identify additional regulators of hydathode formation, we performed a transcriptomic profiling of laser-microdissected, differentiating apical hydathodes from the first *Arabidopsis* leaf pair (we focused on the apical hydathode as its position is highly predictable). Tissues (hydathode versus neighboring regions) were collected over a developmental kinetic starting at 5 days before visible hydathode differentiation, through onset of epithem and water pore differentiation (7 days), until full differentiation (11 days) (Fig. 2A, Extended Data Fig. 6A, 7A, B). These comparisons revealed differentially expressed genes (DEGs) at each time point (Extended Data Fig. 7C, Data S1). The transcriptomic dynamics was confirmed for 8 DEGs by reporter lines (Extended Data Fig. 7D). Pairwise comparisons of hydathode-upregulated DEGs revealed two temporal clusters: an early phase (days 5–7) and a later phase (days 8–11), the latter strongly overlapping with genes previously reported to be expressed in mature hydathodes (Fig. 2A, ^10,30,31^). Gene set enrichment analysis (GSEAs ^32^) showed an under-representation of photosynthesis-related processes at days 8 and 11, while cell-wall related processes (day 8), and inorganic ion transport and bacterium-responsive pathways (day 11) were significantly over-represented (Fig. 2B), consistent with the differentiation and functional specialization of the hydathode ^4,10,33,34^. Together, transcriptomic profiling along with cytological observations pointed to a two-step model of hydathode formation: an early specification phase at days 5–7, followed by a differentiation phase beginning at day 8. Among the DEGs, *STYLISH* 1 (*STY1*) and *STY2* were consistently up regulated in the hydathode domain at all developmental stages, as confirmed by pSTY1:GUS and pSTY2:GUS reporters (Extended Data Fig. 7D, E, 8A). A role for these genes in hydathode formation was further supported by the phenotype of *sty1-1 sty2-1* double mutants that formed leaves with exaggerated teeth and formed no or only small hydathodes at both apical and marginal positions (Fig. 2C-E, Extended Data Fig. 8B).

**Figure 2.**
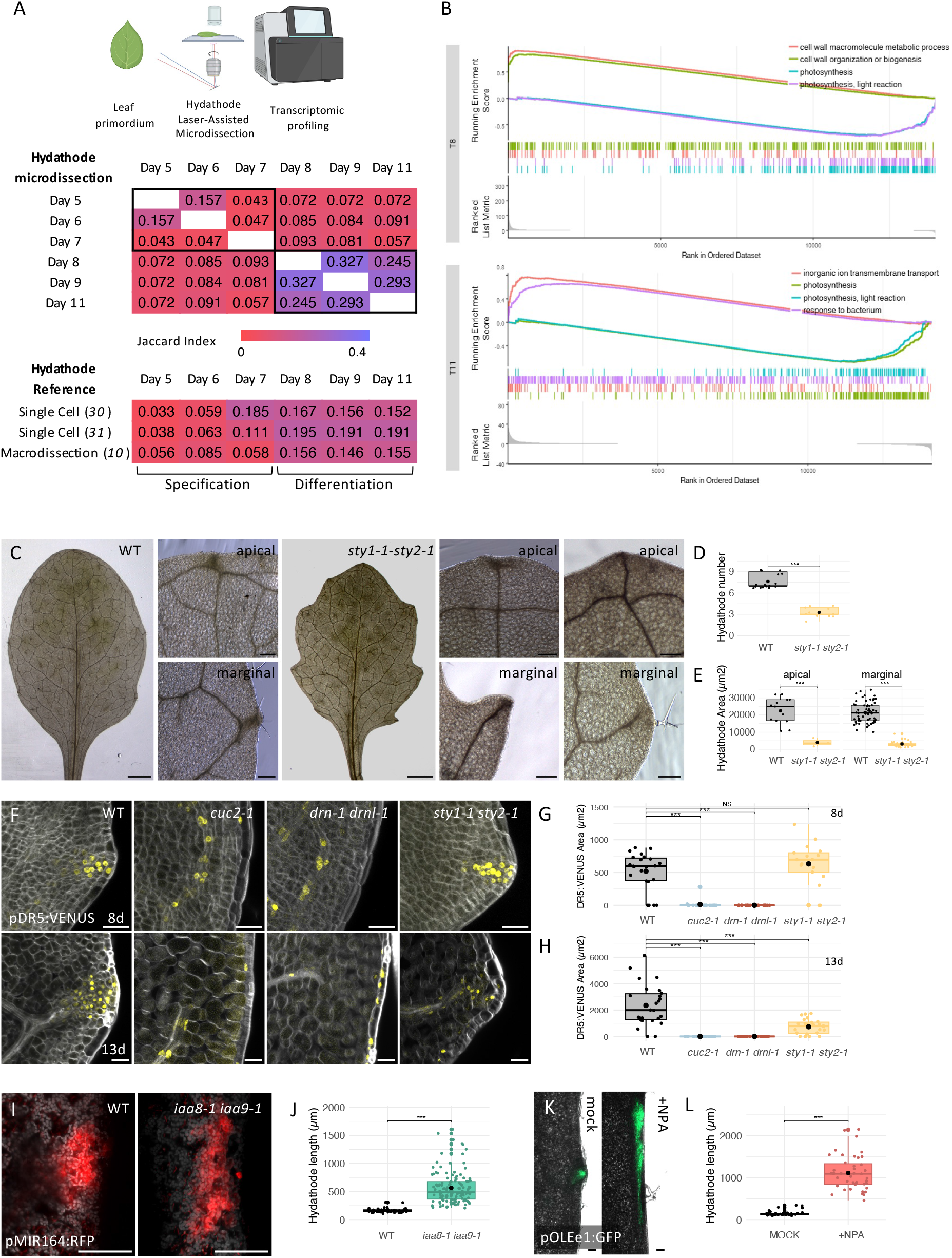
Auxin drives hydathode formation. **A.** Transcriptomic profiling of developing hydathode through Laser-Assisted Microdissection (LAM). Developing hydathodes were isolated by LAM from the first pair of leaves in 5-to 11-day-old seedlings. The first table shows pairwise similarities (Jaccard Index) between genes upregulated in hydathodes at different time points. The second table shows pairwise similarities between genes upregulated in hydathodes at different time points and genes expressed in mature hydathode, as revealed by two single cell transcriptomic profiling ^30,31^ and hydathode macrodissection data ^10^. **B.** Gene Set Enrichment Analysis (GSEA^32^) of DEG at T8 and T11. **C.** Leaf and hydathode phenotype of the *sty1-1 sty2-1* double mutants. For each genotype (WT or *sty1-1 sty2-1*) the first panels show a mature leaf 6, while the smaller upper panels show details of the leaf apex and the lower panels show details of a tooth region. In the mutant, two pictures are shown, with respectively a small hydathode or no visible hydathode. **D.** Quantification of the hydathode number in WT and *sty1-1 sty2-1* double mutants (*n* ≥ 11). **E.** Quantification of the apical and marginal hydathode area in WT and *sty1-1 sty2-1* double mutants (*n* ≥ 3). **F.** pDR5:VENUS expression at the marginal hydathode site at 8 days (upper panels) or 13 days (lower panels), in WT, *cuc2-1* mutants or *drn-1 drnl-1* and *sty1-1 sty2-1* double mutants. **G, H.** Quantification of pDR5:VENUS expression domain area at 8 days (*n* ≥ 17) (**G**) or 13 days (*n* ≥ 23) (**H**), in WT, *cuc2-1* mutants or *drn-1 drnl-1* and *sty1-1 sty2-1* double mutants. **I.** pMIR164A:RFP expression as a marker of hydathode in WT and *iaa8-1 iaa9-1* double mutant. **J.** Quantification of hydathode length (measured along the leaf margin) in WT and *iaa8-1 iaa9-1* double mutant using pMIR164A:RFP expression as a marker of hydathode (*n* ≥ 53). **K.** pOLEe1:GFP-GUS expression as a marker of hydathode in mock- and NPA-treated leaves. **L.** Quantification of hydathode length (measured along the leaf margin) in mock- and NPA treated leaves using pOLEe1:GFP-GUS expression as a marker of hydathode (*n* ≥ 43). Leaves are ethanol-cleared in **C**. VENUS signal is shown in yellow and SR2200 cell wall staining is shown in grey in **F**. pMIR164A:RFP signal is shown in red in **I**, pOLEe1:GFP signal in green in **K** and chlorophyll fluorescence in grey in **I** and **K**. Bars = 2 mm for the entire leaves and 200 µm for the details in **A**, 20 µm in **F** and 100 µm in **I** and **K**.

### Auxin signaling is required and sufficient for hydathode specification

*CUC2*, *DRN*/*DRNL* and *STY* genes pattern plant development by contributing to the formation of spatially-graded auxin responses ^21,27,28,35^. Auxin has previously been proposed to be associated with the formation of the water pores or the epithem ^5,18,20^ and we observed a continuous expression of the auxin signaling reporter pDR5:VENUS at the hydathode sites from specification to differentiation (Extended Data Fig. 9A). Therefore, we tested whether reduction of hydathode development observed in *cuc2-1* mutants and *drn-1 drnl-1* or *sty1-1 sty2-1* double mutants was associated with abnormal auxin response. To do this, we analyzed the expression of the pDR5:VENUS auxin signaling reporter during hydathode specification (at 5 days or 8 days for apical or marginal hydathodes, respectively) and differentiation (at 8 days or 13 days for apical or marginal hydathodes, respectively). In *cuc2-1*, at both stages, pDR5:VENUS expression was normal at the apical hydathode, but was absent from the marginal sites (Fig. 2F-H, Fig. S9B-D). pDR5:VENUS expression was also absent from *drn-1 drnl-1* marginal sites at both stages (Fig. 2F-H). pDR5:VENUS expression was normal during *drn-1 drnl-1* apical hydathode specification, and slightly reduced during its differentiation (Extended Data Fig. 9B-D), in agreement with the smaller size of apical hydathode in this double mutant (Extended Data Fig. 3J). The *sty1-1 sty2-1* double mutant showed normal pDR5:VENUS expression at specification and a strong reduction during differentiation of both apical and marginal hydathodes (Fig. 2F-H, Extended Data Fig. 9B-D). These observations led us to hypothesize that early auxin signaling is required for hydathode specification, whereas sustained auxin signaling is necessary for normal hydathode differentiation. *STY1* induces the expression of *YUC4* ^35^ and *CUC2* promotes the expression of both *YUC1* and *YUC4* genes ^36,37^ that catalyze the last reaction of auxin synthesis in the tryptophan pathways. Accordingly, the *yuc1 yuc4* double mutant developed leaves of normal size with smooth margins that lacked most of the marginal hydathodes (Extended Data Fig. 10). Together, these observations suggest that the formation of hydathodes requires a continuous auxin response that is built up early by the *CUC2* and *DRN*/*DRNL* genes and subsequently maintained by the *STY1*/*STY2* genes, in part through the *YUC* genes. The observation that pDRNL:CER expression is reduced in *cuc2-1* mutants but not in *sty1-1 sty2-1* mutants supports this two-step model (Extended Data Fig. 11).

Next, we wanted to test whether auxin was sufficient to promote hydathode formation. INDOLEACETIC ACID-INDUCED PROTEIN 8 (IAA8) and IAA9 are two auxin signaling repressors that have been proposed to restrict auxin signaling at the leaf margin between outgrowing teeth ^38^. The *iaa8-1 iaa9-1* double mutant formed more hydathodes, sometimes at ectopic positions and that were larger than in the WT (Fig. 2I, J, Extended Data Fig. 12). In WT, treatment with the auxin transport inhibitor N-1-naphthylphthalamic acid (NPA) replaces the regular discrete auxin signaling peaks at the leaf margin by larger domains of auxin response, as revealed by pDR5:VENUS expression (Extended Data Fig. 13A). The hydathodes forming on NPA treated leaves are dramatically enlarged (Fig. 2K, L, Extended Data Fig. 13B-D). Importantly, in both *iaa8-1 iaa9-1* double mutants and NPA-treated plants, the enlarged or ectopic hydathodes showed a coordinated development of epithem, water pores and vascular tissues (Extended Data Fig. 12C, D, 13D, 14). From these observations we concluded that auxin signaling at the leaf margin is both necessary and sufficient for the coordinated development of the different cell types forming tri-partite hydathodes.

### Auxin and cytokinins promote hydathode growth

During leaf development, the size of the epithem domain, marked by E325-GFP expression, is increasing in parallel with the number of epithem cells (Fig. 3A, B). Epithem cells and their nuclei exhibited only a limited increase in size relative to the neighboring spongy mesophyll cells (Fig. 3C, D, Extended Data Fig. 15A-D). No expression of the endoreduplication marker SIAMESE RELATED 1 / LOSS OF GIANT CELLS FROM ORGANS (SMR1/LGO) ^39,40^ was observed in epithem cells, in contrast to clear expression in mesophyll cells (Fig. 3E, Extended Data Fig. 15E). These observations suggest that while spongy mesophyll cells exit mitosis to enter a cell expansion phase associated with endoreduplication, epithem cells remain longer in a proliferative stage. Auxin and cytokinins (CK) promote cell proliferation ^41^, and we observed that treatment of 4-day-old seedlings with 1µM 1-Naphthaleneacetic acid (NAA) auxin or 5 µM 6-Benzylaminopurine (BA) for 4 days led to larger hydathodes (Fig. 3F). CK signaling was revealed in epithem cells by the expression of the TCSn:GFP CK reporter ^42^ (Fig. 3G). Lines overexpressing the negative regulator of CK signaling, RESPONSE REGULATOR 7 (ARR7) ^43^, which expression is induced by DRNL ^44^, showed smaller hydathodes (Fig. 3H, Extended Data Fig. 15F, G). Together, these data suggest that CK signaling positively regulates hydathode growth. CKs act through the regulation of *CYCD3* genes to maintain cells proliferating and restraining them from transition to endoreduplication ^45^. One of the three Arabidopsis *CYCLIN D3* genes, the *CYCD3;1* gene, was specifically expressed in hydathodes (Fig. 3I) and *cycd3;1* mutants showed larger epithem nuclei and larger marginal hydathodes (Fig. 3J, K). Together, these observations support a scenario in which CK signaling in the epithem contributes to restraining the transition to endoreduplication via *CYCD3;1* to maintain cells proliferating and contribute to epithem formation.

**Figure 3.**
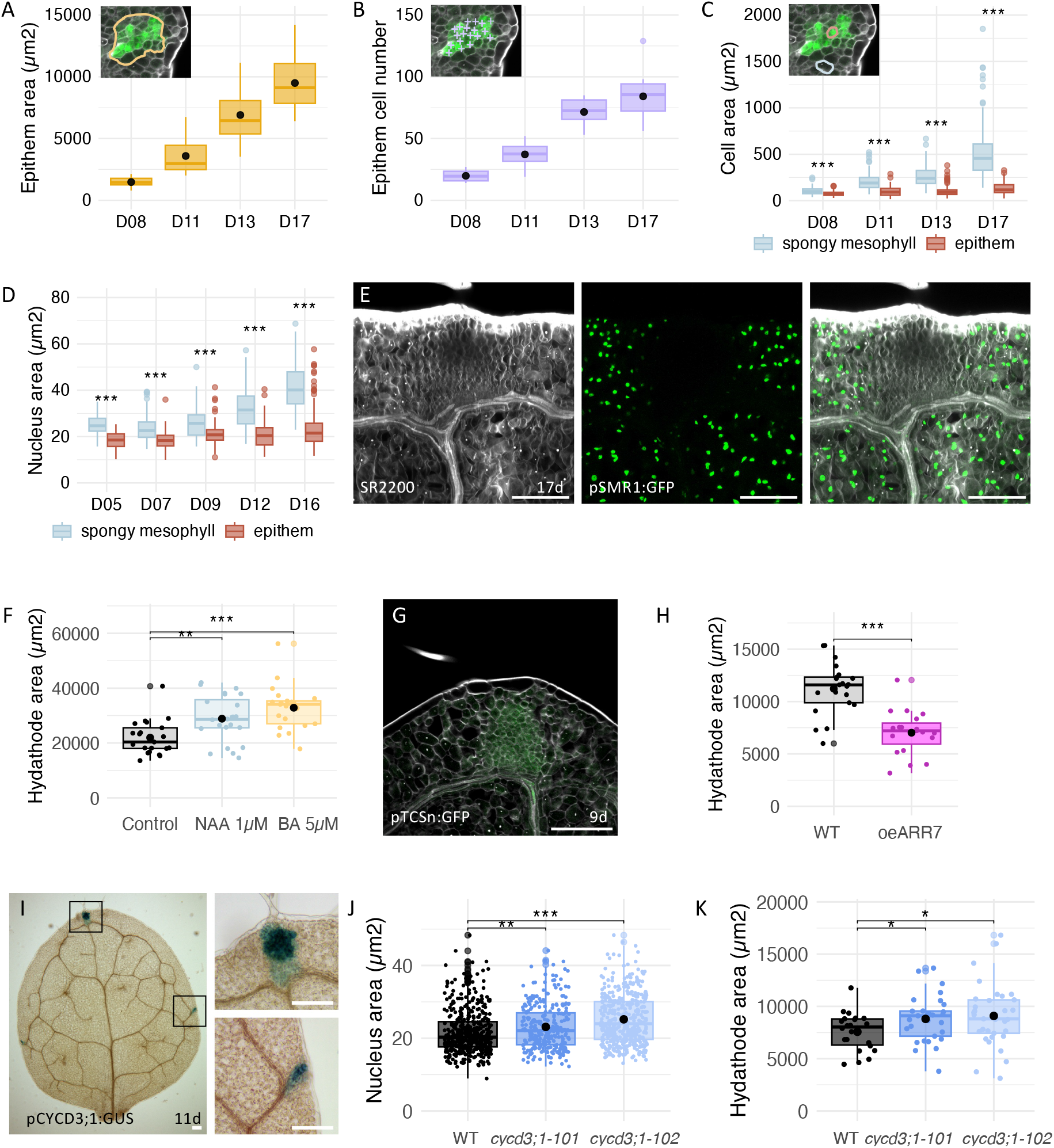
Sustained cell proliferation and limited endoreduplication contribute to epithem formation. **A.** Evolution of epithem size during the development of marginal hydathodes in leaf 1 (*n* ≥ 9). **B.** Evolution of cell number during the development of marginal hydathodes in leaf 1 (*n* ≥ 10). **C.** Evolution of epithem and spongy mesophyll cell sizes during the development of marginal hydathodes in leaf 1 (*n* ≥ 103). **D.** Evolution of epithem and spongy mesophyll nucleus sizes during the development of apical hydathodes in leaf 1 (*n* ≥ 67). **E.** Expression of pSMR1:GFP in the apex (*n* ≥ 10) of leaf 1 at 17 days. **F.** Apical hydathode size (using E325-GFP expression as a proxy) in leaf 1 of 18-day-old seedlings treated with auxin (1 µM NAA) or cytokinin (5 µM BA) for 4 days starting at 4 days after germination (*n* ≥ 21). **G.** Expression of pTCSn:GFP in the apical hydathode of leaf 1 at 9 days. **H.** Marginal hydathode size (using epithem size measured on SR2200-stained sections as a proxy) in leaf 1 of 17-day-old seedlings of the WT and a ARR7 overexpressing line (oeARR7) (*n* ≥ 19). **I.** Expression of pCYCD3,1:GUS in leaf 1 at 11 days. Details show the apical and a marginal hydathode. **J.** Nucleus size in apical hydathodes of leaf 1 at 17 days in the WT and two different mutant alleles of *CYCD3;1*, *cycd3;1-101* and *cycd3;1-102* (*n* ≥ 298). **K.** Marginal hydathode size (using epithem size measured on SR2200-stained sections as a proxy) in leaf 1 of 17-day-old seedlings of the WT and two different mutant alleles of *CYCD3;1*, *cycd3;1-101* and *cycd3;1-102* (*n* ≥ 20). SR2200 cell wall staining is shown in grey and GFP signal is shown in green in **E** and **G**. Bars = 100 µm in **E, G** and **I**.

### Hydathode architecture determines susceptibility to leaf flooding

Next, we took advantage of the identification of mutants with shared altered hydathode formation but contrasted effects on overall leaf development to perform a genetic analysis of hydathode function. When exposed to conditions that induce guttation in the wild type such as elevated air humidity (Extended Data Fig. 16A), the five lines with fewer and/or smaller hydathodes (*cuc2-1*, *drn-1 drnl-1*, *drn-3 drnl-cr4, sty1-1 sty2-1*, and *oeARR7*) consistently showed leaf flooding, while the WT and mutants with larger hydathodes (*iaa8-1 iaa9-1* and *cycD3-101, cycD3-102*) showed either no or limited flooding (Fig. 4A, B, Extended Data Fig. 4G, 16B). Flooded areas appeared as dark patches that were readily detected using a short wave infra-red imaging (Extended Data Fig. 17A-B). While mechanical blockage of hydathodes in *Chloranthus japonicus* led to flooding restricted to the leaf margin ^14^, genetic disruption of hydathode function led to flooded sectors of variable size with no preferential localization in the leaf blade (Fig. 4A). The extent of leaf flooding was dependent on genetic background, as the same *cuc2-1* mutant allele led to more severe flooding in the L*er* background than in Col-0 (Extended Data Fig. 16C). Leaf flooding is also developmentally regulated, as leaves of the same rank showed different flooding intensities depending on their developmental stage (Extended Data Fig. 16D). When plants were removed from the conditions inducing guttation (lowering humidity), leaf flooding rapidly disappeared, with a faster rate of recovery observed in *cuc2-1* than in *sty1-1 sty2-1* (t_50_ half time to full recovery of 16±4 min and 43±7min, mean±SE respectively) (Fig. 4C, Extended Data Fig. 17D, E, Movie1-2). This rate of recovery was faster when plants were exposed to environments with lower atmospheric humidity (Fig. 4D). Furthermore, when repetitively exposed to daily cycles of guttation-inducing conditions, flooding consistently reappeared in the overlapping leaf regions, suggesting that it may initiate from structural heterogeneities within the leaf, such as weak points along the vascular network (Extended Data Fig. 16E). Altogether, these observations indicate that hydathode size and number are critical for preventing leaf flooding, a process that is reversible and regulated by both developmental and environmental cues.

**Figure 4.**
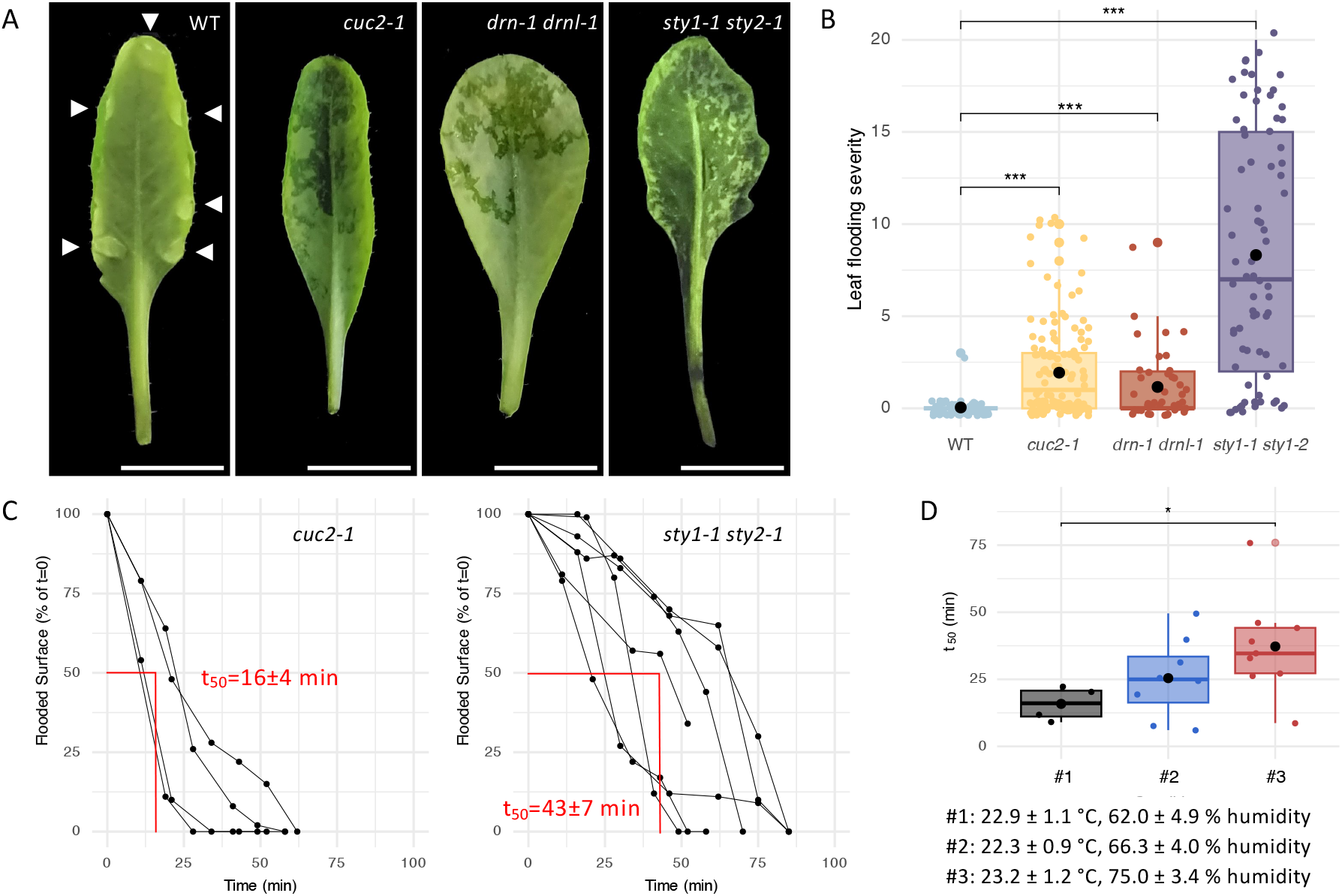
Hydathodes are necessary to limit reversible, environmentally-controlled leaf flooding. **A.** Leaves following over-night induction of guttation show guttation droplets (indicated by white arrowheads) or leaf flooding (visible as dark patches on the leaf blade). The leaf lower side of WT, *cuc2-1* mutants, *drn1 drnl-1* and *sty1-1 sty1-2* double mutants are shown. **B.** Quantification of leaf flooding severity in WT, *cuc2-1* mutants, *drn1 drnl-1* and *sty1-1 sty1-2* double mutants. Leaf flooding severity is indicated by a score ranging from 0 (no flooding) to 20 (100% of the leaf blade area flooded) (*n* ≥ 44). **C.** Kinetic of leaf flooding reversion in *cuc2-1* mutants and *sty1-1 sty1-2* double mutants. The evolution of the flooded area (normalized to the area flooded at t=0) is indicated. Measures at different moments (points) of the same leaf are connected by thin black lines. The mean±SE t_50_ (necessary for recovering half of the starting flooded area) is indicated. **D.** t_50_ values measured for the reversion of leaf flooding in *cuc2-1* mutants under 3 different environmental conditions (#1 to #3). Details of the conditions are in indicated below the graph. Condition #1 was used in C (*n* ≥ 4).

## Discussion

Here, we provide a detailed cellular, genetic and molecular framework for hydathode formation in *Arabidopsis* (Fig. 5). This framework comprises two interconnected regulatory modules acting sequentially. The first module, which leads to the specification of hydathodes, relies on auxin signaling that is both necessary and sufficient for the coordinated development of the three cell types forming hydathodes, xylem, epithem, and water pores. The second module involves the combined action of auxin and CK, which act together to maintain cell proliferation and delay endoreduplication, thereby contributing to the formation of small epithem cells. The functional importance of maintaining epithem cells small, particularly in the context of the enlarged mesophyll cells, remains to be explored. Importantly, our findings provide a molecular basis for the spatial and temporal coordination of these two modules. Auxin, required throughout all stages of hydathode formation, likely ensures spatial coordination. We further propose that temporal coordination of the two modules occurs through the *DRN*/*DRNL* genes. Because these genes have been shown in other organs to activate *STY* expression and to repress CK signaling ^44,46–48^, we propose that their early transient expression in developing hydathodes determines the timing of the transition from the specification phase to subsequent proliferation and differentiation phases. Although our data suggest that the regulators analyzed here promote hydathode development primarily through the auxin pathway, they may also have more direct effects on hydathode differentiation. For example, *STY* genes, which promote auxin synthesis via the activation of *YUC* genes, have also been proposed to activate the cell-wall–modifying genes *POLYGALACTURONASE-LIKE 1* (*PGL1*) and *EXPANSIN-LIKE A2* (*EXLA2*), two genes expressed in developing and mature hydathodes ^10,49^ (Extended Data Fig. 7F).

**Figure 5.**
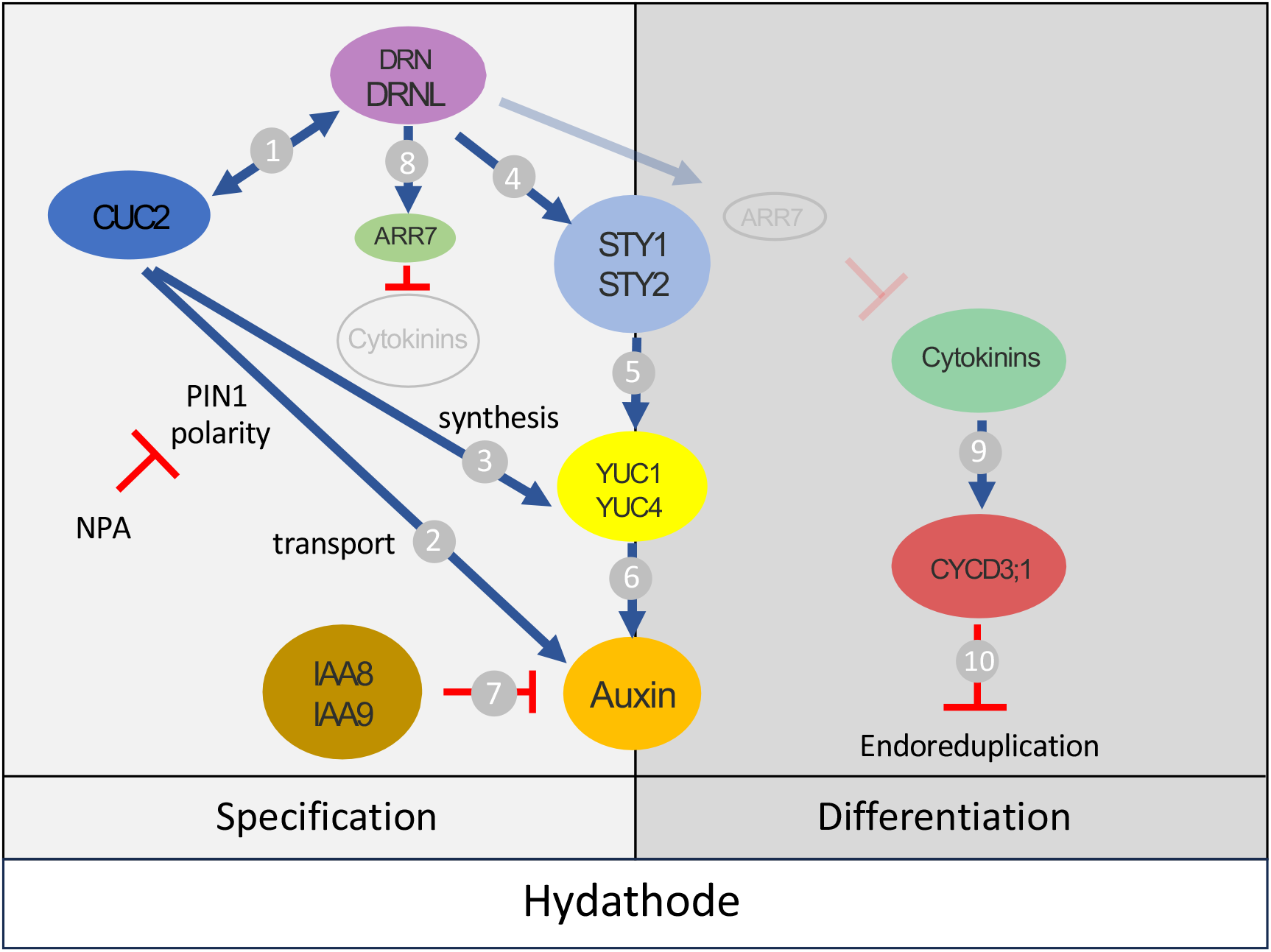
Model of hydathode formation in Arabidopsis. Specification of the hydathodes along the leaf margin requires the interacting *CUC2* and *DRN*/*DRNL* genes, which contribute to local auxin response either by polar auxin transport or by activating local auxin synthesis through the STY1/STY2/YUC1/YUC4 pathway. Activities of *IAA8* and *IAA9* genes inhibit auxin response. Cytokinin response is delayed via the activity of the *DRNL* gene. This general model is for marginal hydathodes. *DRN* is not expressed in the apical hydathodes. *CUC2* is not required for apical hydathodes and we propose that *CUC1* and/or *CUC3* may have redundant roles, as the auxin response at the leaf primordium is already set up in the meristem when the leaf primordium is initiated. Auxin and cytokinins contribute to hydathode (epithem) proliferation and limit endoreduplication. Cytokinins are acting through the activation of the *CYCD3,1* gene which represses endoreduplication. Support for the different interactions and contribution to hydathode formation are the following. **1.** Genetic interactions between *CUC2*/*DRN*/*DRNL* in different organs ^26–28^. This study: absence of *DRNL* expression in *cuc2* mutant margins. **2.** *CUC2* promotes auxin response through polar auxin transport via the PIN1 proteins ^21,22,60^. This study: *cuc2* hydathode defects, reduced auxin response at hydathode sites in *cuc2*, NPA effects on hydathode formation. **3.** *YUC1* and *YUC4* expression is activated by *CUC2* ^36,37^. **4.** *DRNL* promotes the expression of *STY1* and other members of the family ^48^ This study: *drn* and *drn drnl* hydathode defects, reduced auxin response at hydathode sites in *drn drnl*, *DRN*/*DRNL* expression patters, *DRNL* expression does not require *STY1*/*STY2*. **5.** *STY* promotes *YUC* expression ^35^. This study: *sty1 sty2* hydathode defects, reduced auxin response at hydathode sites in *sty1 sty2*, *STY* and other family member expression patterns. **6.** YUC encode auxin biosynthesis genes ^61^. This study: *yuc1 yuc4* hydathode defects. **7.** *IAA8* and *IAA9* repress auxin signaling during leaf development ^38^. This study: *iaa8 iaa9* hydathode defects. **8.** *DRNL* promotes *ARR7* expression and repress cytokinin signaling signaling ^44,46,47^. **9.** Cytokinins induce *CYCD3* expression ^62^. This study: TCSn:GFP expression, effects of cytokinins on hydathode phenotype, OEARR7 hydathode phenotype. **10.** *CYCD3,1* represses endoreduplication ^45,63^. This study: *CYCD3,1* expression and *cycD3,1* hydathode and endoreduplication phenotype.

Together, our findings also reveal how hydathode formation is integrated into the broader leaf developmental program. Auxin distribution and signaling, which are central for hydathode formation, also determine tooth and vascular patterning ^21,50–52^, thereby ensuring the functional continuity of these two structures with hydathodes. The spatial connection of hydathodes with the vasculature ensures hydraulic continuity, while their positioning at leaf and tooth tips may optimize guttation droplet removal. However, our results also demonstrate that hydathode and tooth formation can be uncoupled, suggesting that these processes differ in their timing or sensitivity towards auxin signaling. Beyond auxin, the genetic factors identified as key regulators of hydathode formation also impact leaf morphogenesis and are part of the larger network determining overall leaf size, shape, or serration pattern ^53–55^. Our work therefore reveals how the redeployment of common developmental regulators enables the emergence of spatially restricted structures endowed with specialized physiological functions within a growing organ. Similarly, in mammalian tooth development a conserved gene regulatory network is deployed iteratively, first establishing the overall tooth pattern and subsequently generating cusps ^56,57^. Importantly, our results also provide hypotheses for how hydathode formation and leaf morphogenesis can become uncoupled, opening evolutionary scenarios for the diversification of hydathode number and position in plants. Finally, by identifying mutants impaired in hydathode formation, our study establishes a foundation for the genetic dissection of hydathode function in plant physiology, as illustrated here by their role in the prevention of leaf flooding.

## Acknowledgements

We thank Y Shi (China Agricultural University, Beijing), W. Werr (University of Cologne, Germany), J Murray (University of Cardiff, Great Britain), M Aida (Kumamoto University, Japan), K Torii (Nagoya University, Japan), C Galvan-Ampudia (ENS de Lyon, France) and the NASC for providing seeds, and N Arnaud, G Venkatesh, M Caroulle for providing comments on the manuscript.

## Author Contributions

Conceptualization: PS, LN, NL, PL Methodology: NR, NL, BG

Investigation: PS, ASS, PP, DV, AO, SP, CPLR, CQ, NL, PL

Resources: JMR

Writing – original draft preparation: PS, PL

Writing – review and editing: all authors

Supervision: PS, NL, PL

Funding acquisition: LN, NL, PL

## Data and materials availability

The RNA-Seq data were deposited at NCBI/GEO (^58^, http://www.ncbi.nlm.nih.gov/geo/) with accession number GSE308406. All steps of the experiment, from growth conditions to bioinformatic analyses, were detailed in CATdb database (^59^, http://tools.ips2.u-psud.fr.fr/CATdb/) under project name 2023_03_QS_cinedathodes. All other data are available in the main text or supplementary materials. The described biological material can be obtained from PL.

