## Supplemental for "Genetic coupling of hydathode formation with leaf morphogenesis maintains water homeostasis"

Pauline Savourat et al

**The PDF file includes:**

Materials and Methods
Extended Data Figs. 1 to 17
Extended Data References
Extended Data Table 1

**Other Supplementary Material for this manuscript includes the following:**

Extended Data Set 1
Extended Data Movie 1 and 2

### Material and Methods

**Plant Material**

The following lines were described before: e325-GFP [1], pOLEe1:GFP-GUS, pPGL1:GFP-GUS [2], *cuc2-1* (back-crossed into Col-0) [3] *cuc2-1* (Ler) [4] *cuc2-3* pCUC2:CUC2:VENUS [5], pMIR164A:erRFP, pDR5:N7-VENUS [6], *drn-2, drn-1 drnl-1* [7], pDRNL:erCER [8], pDRN:erGFP [9], *sty1-1 sty2-1*, pSTY1:GUS, pSTY2:GUS [10], pSHI:GUS [11], pSRS5:GUS [12], pSRS7:GUS, pLRP1:GUS [13], *yuc1 yuc4* [14], *iaa8-1 iaa9-1* [15] pCYCD3;1:GUS [16], oeARR7 [17], pSMR1::nlsGFP-GUS [18], pTCSn:GFP [19]. The *drn-3* (SALK_030262), cycd3;1-101 (SM_3.24877) and cycd3;1-102 (GK-529C07) lines were obtained from the Nottingham Arabidopsis Stock Centre.

**Plant growth**

Plants were grown *in vitro* on Arabidopsis medium (Duchefa, prod. n° DU0742.0025) added with 0,33 g.L$^{-1}$ $Ca(NO_3)_2$ and 20 mM MES, pH 5.8 under long day conditions [16h light / 8h dark at 21°C] after 48h vernalization at 4°C. Plants were grown in a greenhouse, under long-day conditions.

NPA treatment was performed as described in [6]

**Genome editing of *DRNL*.**

CRISPR-Cas9-mediated genome editing was used to generate *drnl* mutants in Col-0 background. Briefly, two RNA guides (gRNA1: TGCGCCGCTCGAGCCATGCG and gRNA2: GATACGGGACCCATTGTCCA) were designed to target the AP2 region of *DRNL* and two RNA guides to the C-terminal domain (gRNA3: GACATCGTTGACGTTAGTGA and gRNA4: GAACCAGAACCAGCTAGTTC). Using the GoldenBraid method [20] the gRNAs were placed under the control of the U6 promoter, combined two by two (gRNA1 + gRNA2 and gRNA3 + gRNA4) and finally combined with a pCMV:DSRed:tnos pRPS5a:hcas9:TRbcSE9 module. The two resulting constructs were introduced into wild-type Col-0 via *Agrobacterium*-mediated transformation using a floral dip method. Primary transformants were identified using DSRed expression in the seeds as a marker. T2 plants having segregated away the transgene were identified by the absence of DSRed expression in the seeds and the *DRNL* locus was sequenced. Three plants homozygous for mutations in *DRNL* were identified (*drnl-cr2*, resulting from cleavage at gRNA3 site; *drnl-cr3*, resulting from cleavage at gRNA2 site; *drnl-cr4*, resulting from cleavage at gRNA1 and gRNA2 sites). The *drnl-cr4* mutant was crossed with *drn-3* and *drnl-cr4 drn-3* double homozygous plants were identified in the F2 generation. However, because these *drnl-cr4 drn-3* double mutants were sterile, we also identified fertile F2 plants homozygous for *drnl-cr4* and heterozygous for *drn-3* which generated one quarter of *drnl-cr4 drn-3* double mutants in their progeny.

#### Imaging

##### *Sample preparation*

For confocal imaging, samples were fixed under vacuum on ice with 4% (w/v) PFA, 0.1% Triton in 1X PBS for 1 hour and, after renewal of the fixative, were left over night at 4°C. Fixed tissues were washed twice in 1X PBS and cleared with ClearSee [21] at room temperature for several days. Cell wall staining was performed by replacing the initial ClearSee solution by 0.1% SCRI Renaissance 2200 (SR2200) dye in ClearSee for at least one day before imaging. For DNA staining using DAPI, cleared samples were washed twice in 1X PBS, stained 10 minutes in 1 $\mu g.ml^{-1}$ DAPI and observed after two washes in 1X PBS.

To count hydathodes, leaves were cleared in ethanol. For this, leaves were immersed in 70% ethanol, heated in successive cycles of approximately 20 seconds each, until boiling was reached. The process was repeated until sufficient clearing was obtained, with renewal of the ethanol solution. After clearing, the leaves were rehydrated progressively and were kept and observed in a 30% glycerol solution.

The GUS staining was done as described by [22] in the presence of 2mM of potassium ferrocyanide and 2 mM of potassium ferricyanide. The emergence of the blue staining was regularly checked and the staining was stopped when clearly visible by washing the samples twice in water and clearing them in 70% ethanol. The same staining duration was applied to different genotypes carrying the same reporter.

*Confocal microscopy and macroscope imaging*

Confocal imaging was performed on a Leica SP5 inverted microscope for hydathode structure and gene expression characterization or on Zeiss 710 for DAPI imaging. Acquisition parameters are presented in Extended Data Table 1. Maximum intensity projections along the Z axis were made using ImageJ. When necessary, stitching of several pictures was performed using the Pairwise Stiching plugin [23] and FigureJ [24] was used to assemble pictures into figures.

GUS staining or larger leaves expressing fluorescent reporters were observed using an Axio Zoom.V16 macroscope (Carl Zeiss Microscopy, Jena, Germany, http://www.zeiss.com/)

*SWIR Imaging*

Leaf flooding was monitored using short-wave infrared (SWIR) imaging, which allows non-invasive visualization of water accumulation in plant tissues based on the strong absorption of SWIR radiation by liquid water. Plants were imaged using an InGaAs-based SWIR camera sensitive in the 900–1700 nm spectral range. Illumination was provided by led light source positioned at a fixed angle relative to the leaf surface to ensure homogeneous illumination and minimize reflections. Plants were imaged *in situ* under controlled environmental conditions. For flooding assays, plants were exposed to high relative humidity (100%) to induce guttation. Leaves were imaged at defined time points during exposure to high humidity and following the return to control conditions (66% relative humidity). Camera exposure time, gain, and working distance were kept constant across all genotypes and treatments. Images were acquired in reflectance mode. Flooded leaf areas appeared as darker regions in SWIR images due to increased absorption of infrared radiation by water. For visualization and quantification, images were processed using ImageJ software. Pixel intensity values were normalized to background and, when required, thresholded to segment flooded regions. Flooded area was expressed as a proportion of total leaf area.

**Image Quantification**

The number of hydathodes was counted manually on ethanol-cleared leaves or using hydathode markers.

Hydathode area was measured manually with FIJI [25], using E325-GFP or pOLEe1:GFP as markers in confocal pictures for *cuc2-1*, *drn-1 drnl-1* and WT.

Hydathode area was measured manually with FIJI based on the small size of epithem cells on SR2200 stained confocal pictures for *oeARR7*, *cycd3;1-101* and *cycd3;1-102* mutants and *sty1-1 sty2-1* double mutants.

Hydathode length was measured manually with FIJI, using pMIR164A:RFP expression as a marker in *iaa8-1 iaa9-1* and WT on confocal pictures, or using E325-GFP or pOLEe1:GFP-GUS as a marker in NPA-treated or control plants on macroscope pictures.

The size of the hydathode after NAA or BA treatments or control plants was measured using the Q-pix FIJI macro [5,26] using E325-FGP as a marker on macroscope pictures.

pDR5:VENUS and pDRNL:CER expression domain area was measured manually on Z-stack maximum projections using FIJI.

Nucleus area was measured with FIJI by drawing a ROI on confocal optical section of DAPI-stained tissues. To measure the nucleus volume and the DAPI signal, confocal stacks were first analysed with PlantSeg [27] using a 3D root nuclei model and next 3D nuclei were reconstructed with FIJI using the "3D Objects Counter" plugin [28] and their volume and DAPI signal were measured using the "3D Image J suite" [29].

**Statistical analysis and data visualization.**

All data were analysed using R [30]. Boxplot were generated using the geom-boxplot function of the ggplot2 package [31], combined with individual measures and average value (shown as a black point). These boxplots compactly display the distribution of a continuous variable by allowing the visualization of the median (bold horizontal bar), the first and third quartile (respectively lower and upper hinges), the lowest and largest values no further than 1.5 * interquartile range from the hinge (respectively lower and upper whisker) and outliers (displayed as individual points). Statistical significance is tested by Student tests: NS for not significant, p-value is <0.05 for *, <0.01 for **, <0.005 for ***.

**Laser-Assisted Microdissection and RNA extraction**

Freshly sampled leaves from in vitro grown seedlings were microdissected with the ZEISS PALM MicroBeam using the Fluar 5x/0.25 M27 objective using MMI membrane slides (Prod. No. 50103) and microdissected samples collected in ZEISS AdhesiveCaps. Approximately 10 microdissected tissues (hydathode versus neighboring regions) were collected for each replicate (3 replicates per time point). Total RNAs were extracted using the Arcturus PicoPure RNA Isolation (Applied Biosystems, Prd. n° 15295033) kit following manufacturer's instruction with an additional step with RNase-free DNase set (Prod. N° 79254 QIAgen). RNA quality was controlled using the Agilent RNA 6000 Pico Kit.

**Transcriptomic profiling**

*Library construction*

RNA-seq libraries were constructed with 1ng of total RNA using the Lexogen QuantSEQ 3' mRNA-Seq Library Prep Kit, ref 015.96 (96 rxn) with UMI according to the supplier's instructions for low quantity. Libraries quality were checked on Agilent DNA HS chip, ref #5067-4627 and were quantified with Quant-iT™ PicoGreen™ dsDNA Reagent, ref #P7581 (Thermo Fisher Scientific Inc) to generate an equimolar pool. Libraries were sequenced in single-end mode with 75 bases for each read on a NextSeq500 (Illumina).

*Preprocessing data*

Read 2s were excluded from the pre-processing. UMIs were removed and appended to the read identifier with the extract command of UMI-tools (v1.0.1, [32]). Reads where any UMI base quality score falls below 10 were removed. To remove adapter sequences, poly(A), poly(G) sequences, and low-quality nucleotides, reads were trimmed with BBduk from the BBmap suite (v38.84, [33]) with the options k=13 ktrim=r useshortkmers=t mink=5 qtrim=r trimq=10 minlength=30. Trimmed reads were then mapped using STAR (v2.7.3a, [34]), with the following parameters --alignIntronMin 5 --alignIntronMax 60000 --alignMatesGapMax 6000 –alignEndsType Local--outFilterMultimapNmax 20 --outFilterMultimapScoreRange 0 –outSAMprimaryFlag AllBestScore –mismatchNoverLmax 0,6 on the *Arabidopsis thaliana* genome reference (TAIR www.arabidopsis.org). Reads with identical mapping coordinates and UMI sequences were collapsed to remove PCR duplicates using the dedup command of UMI-tools with the default directional method parameter. RSeQC (v2.6.6, [35]) was used to evaluate deduplicated mapped reads distribution. Deduplicated reads were counted using HTSeq version v0.12.4 [36] (htseq-count mode intersection-nonempty) based on the gene annotations. Between 0,1 (45,3% of raw reads) and 1,8 (79,3% of raw reads) millions of deduplicated reads were associated to annotated genes.

*Genome and annotations*

Genome and gene annotations, including the GO and KEGG if available, come from the EnsemblPlant database (https://plants.ensembl.org/)

*Statistical Analyses of Expression Data*

Statistical analyses were conducted on R v3.6.2. [37] using the R script-based tool DiCoExpress [38,39] based on the Bioconductor package edgeR (v 3.28.0 [40,41]). Hydathode domains and its control domains were compared along the kinetics by comparing (i) T5-T6, (ii) T6-T7, (iii) T7-T8, (iv) T8-T9 and (iiv) T9-T11.

For each analysis, genes with low counts were filtered using the “filterByExpr” function where the group argument specifies the biological conditions, the min.count value is set to 2 and the other arguments are set to their default values. Libraries were normalized with TMM method with the default parameter values.

Each differential analysis was based on a negative binomial generalized linear model in which the logarithm of the average gene expression is an additive function of times, area, their interaction, and a replicate effect. Based on the experimental design, DiCoExpress automatically generated the following contrasts: the difference between two consecutive time points for each area and averaged on the two areas, the difference between the two areas at each time point and averaged on two time points and finally the biological interaction defined as the difference between the two areas at the first time point minus the difference between the two areas at the second time point. Only biologically meaningful contrasts were interpreted. For each contrast a likelihood ratio test was applied and raw p-values were adjusted with the Benjamini–Hochberg procedure to control the false discovery rate We checked that the distribution of the raw p-values followed the quality criterion described by Rigaill et al., 2018 [42] and declared that a gene was declared differentially expressed if its adjusted p-value was lower than 0.05. If the sign of its log fold-change is positive, then the gene is declared overexpressed. If the sign of its log fold-change is negative, then the gene is declared underexpressed. Gene list overlaps and Jaccard index were assessed using the R package GeneOverlap (v1.44.0; [43]) based on a hypergeometric test, with p-values adjusted for multiple testing using the Benjamini–Hochberg method (FDR < 0.05). Gene Set Enrichment Analysis (GSEA) was performed using clusterProfiler (v4.16.0; [44]), and enriched terms were considered significant at FDR < 0.05.

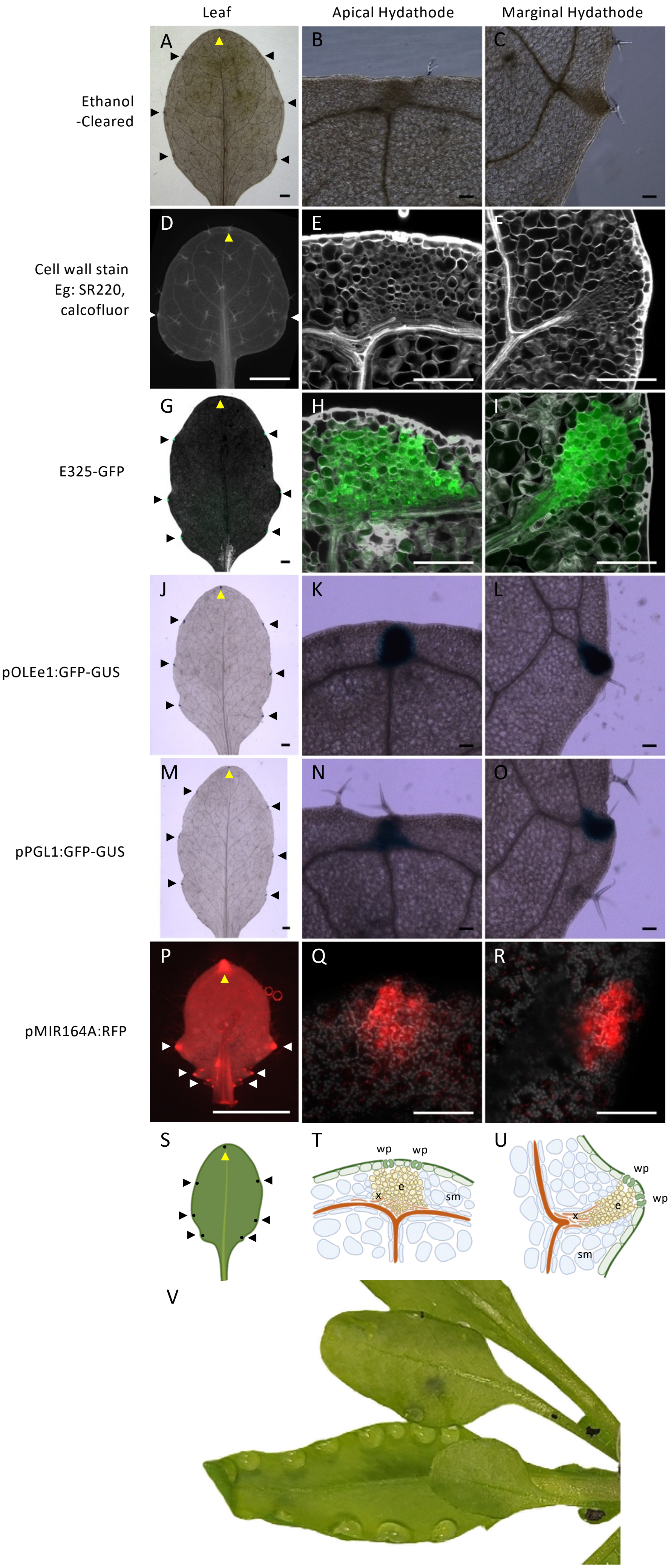


Extended Data Figure 1

Savourat et al.

**Extended Data Figure 1: Multiple molecular, cellular and functional markers allow hydathode identification.**
In ethanol cleared leaves (**A-C**), hydathodes appear as dense structures located close to the leaf margin and connected to the vascular network.
Following cell wall staining by calcofluor or SR2220 and clearing by ClearSee [21] (**D-F**), the small cells forming the epithem can be recognized within the larger spongy mesophyll cells. Xylem strands connected to the epithem are visible. Water pores can also be recognized (See Extended Data Figure 2).
Several reporter lines are expressed in the epithem cells, such as the E325-GFP enhancer trap line [1] (**G-I**), the pOLEe1:GFP-GUS [2] (**J-L**) and pPGL1:GFP:GUS [2] (**M-O**) reporters or the pMIR164A:RFP reporter [6] (**P-R**). Note pMIR164A:RFP is also expressed in the young sinuses between outgrowing teeth, an expression that disappears when the teeth are becoming bigger.
Hydathodes are located around the leaf and an apical hydathode located at the distal leaf tip can be distinguished from marginal hydathodes located along both leaf sides (**S**). The schematic representation of the hydathode shows the organization of the tissues in the apical and marginal hydathode (**T**, **U**). The cells of the epithem (yellow) are smaller than the mesophyll cells (blue); the conducting vessels (orange) and the tracheids (light orange) supply the epithem. The epithem is connected to pores that resemble large stomata located in the epidermal layer (green). Hydathodes can also be recognized functionally by the guttation they mediate and which is visible at the lower side of the leaves (**V**).
Yellow arrow heads point to apical hydathodes and black/white arrow heads point to marginal hydathodes in (**A**, **D**, **G**, **J**, **M**, **P, S**). Mature leaf 6 (**A**, **J, M**), leaf 1 (**D),** leaf 9 (**G**) or developing leaf 11 (**P**) are shown. Samples were imaged using a macroscope (**A**-**C**, **J**-**O**) or a confocal microscope (**D-I**, **P-R**). SR220 or calcofluor staining are shown in grey in (**E-F**) and (**H-I**), respectively, while chlorophyl fluorescence in shown in (**Q-R**). Bars= 1mm for the leaves and 100 µm for the hydathode details in all panels. In **T** and **U**, e: epithem; wp: water pore; sm: spongy mesophyll, x: xylem.

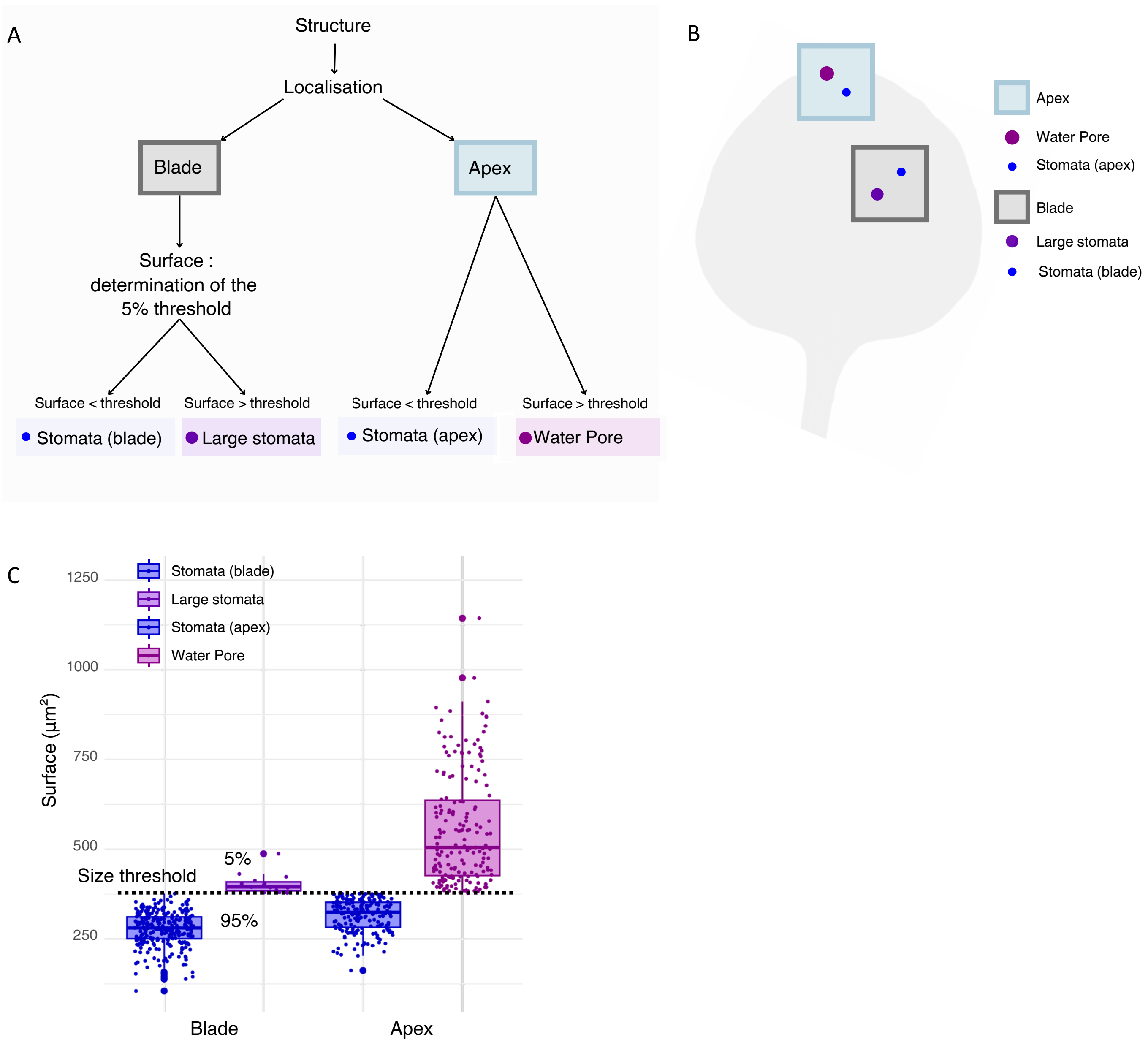


**Extended Data Figure 2: Identification of water pores based on their size and localization.**
Water pores (also called hydathode pores) are stomata-like structures located in hydathodes close the epithem and which in many species, like in *Arabidopsis thaliana* are larger than stomata [45,46]. We took these two criteria, i) localization next the an hydathode (epithem) and ii) large size to identify water pores (**A, B**). First, we analyzed the size distribution of true stomata measured at the leaf blade and defined a size threshold which corresponded to 5% of the largest stomata. This defined two subsets within the stomata population, stomata, and large stomata cells (**C**). Next, we measured the size of stomata-like structures at the leaf apex, around the apical hydathode, which is expected to contain water pores and stomata. We applied the size threshold defined above (represented by a horizontal dotted line) to identify water pores from stomata (**C**).

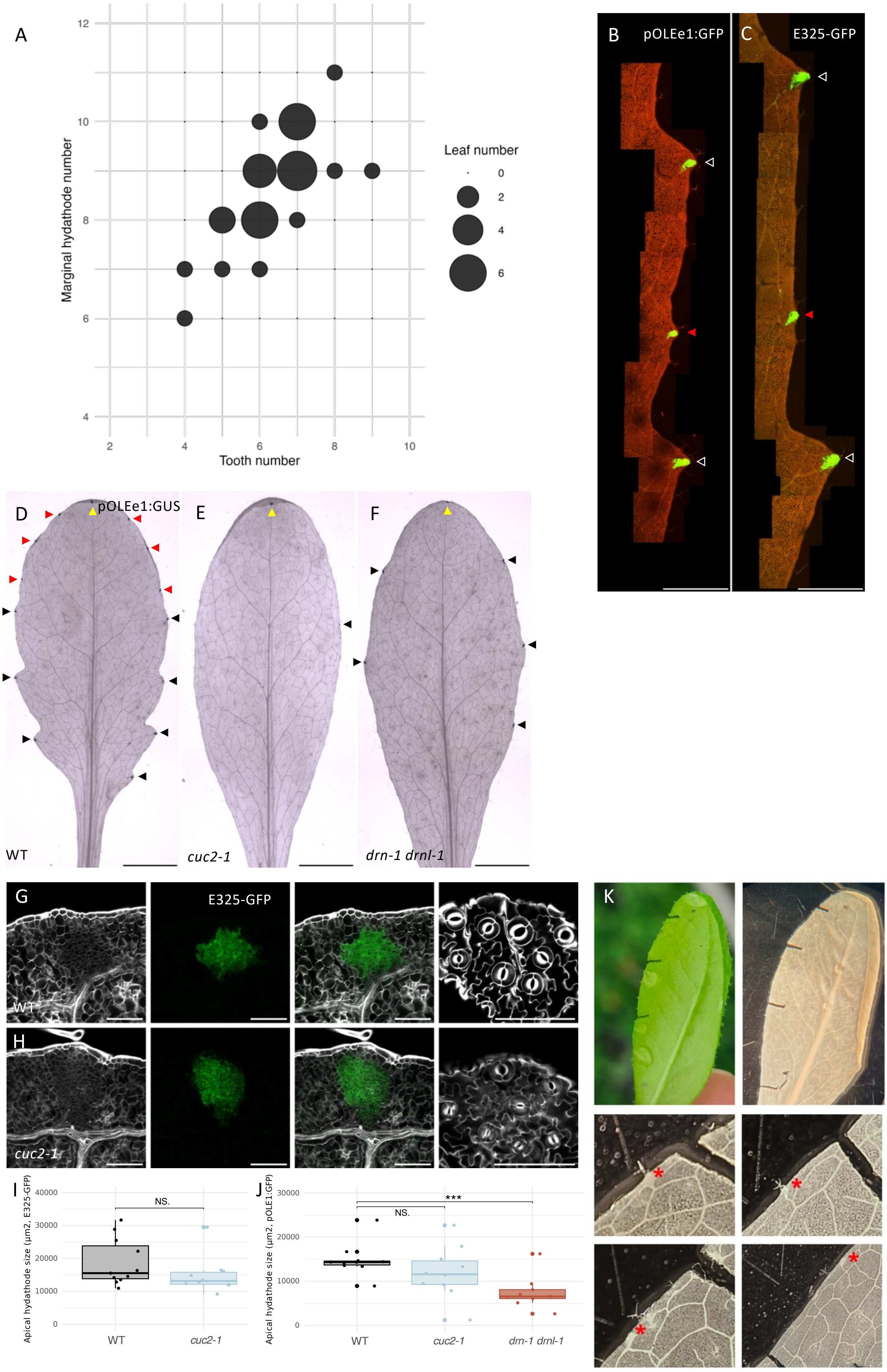

A
Marginal hydathode number
Tooth number
Leaf number
0
2
4
6
B
pOLEe1:GFP
C
E325-GFP
D
pOLEe1:GUS
E
F
WT
cuc2-1
drn-1 drnl-1
G
E325-GFP
WT
H
cuc2-1
K
I
Apical hydathode size (µm2, E325-GFP)
NS.
WT
cuc2-1
J
Apical hydathode size (µm2, pOLE1:GFP)
NS.
***
WT
cuc2-1
drn-1 drnl-1

**Extended Data Figure 3. Hydathodes can be uncoupled from tooth fate.**

**A.** The number of marginal hydathodes is not strictly correlated to the number of teeth. Sizes of the circles indicate the number of leaves showing each combination of leaf teeth / marginal hydathodes (*n* = 36).

**B, C.** pOLEe1:GFP-GUS and E325-GFP expression as markers of hydathodes in a part of a wild-type leaf, showing two hydathodes located each in a tooth and a central hydathode independent of any morphologically-visible tooth.

**D, E, F.** pOLEe1:GFP-GUS expression as a marker of hydathodes in a WT, *cuc2-1* and in *drn-1 drnl-1* leaf 11. Although the number of hydathodes is reduced in both mutants, they still form a few marginal hydathodes.

**G, H.** E325:GFP and SR2200 cell staining showing a similar organization of the apical hydathode in WT and *cuc2-1*. The three first panels show a section through the epithem while the last one shows the water pores.

**I.** Leaf 1 apical hydathode size (using E325-GFP expression domain area as a proxy) is similar in WT and *cuc2-1* ($n \geq 10$).

**J.** Leaf 1 apical hydathode size (using pOLEe1:GFP-GUS (GFP) expression domain area as a proxy) is similar in WT and *cuc2-1*, but reduced in *drn-1 drnl-1* compared to WT ($n \geq 9$).

**K.** Hydathodes formed in *cuc2-1* can mediate guttation. The first panel shows guttation on the lower side of a *cuc2-1* leaf. Small marks are made along the leaf margin to record the position of the guttation droplets. The second panel shows the same leaf which was ethanol-cleared. The four following small panels show details of the leaf margin, with visible hydathodes (red asterisks) which positions are close to the ones of the guttation droplets of the first panel.

Yellow arrow heads point to apical hydathodes and black arrow heads point to marginal hydathodes associated with a tooth while red arrow heads point hydathodes not a associated with a tooth in **B-F**. GFP signal is shown in green in **B**, **C**, **G** and **H**, fluorescence of the chlorophyl is shown in red in **B** and **C**, and grey shows SR2020 cell wall staining in **G** and **H.** Bars= 1 mm in **B** and **C**, 5 mm in **D**-**F**, and 100 µm in **G** and **H**.

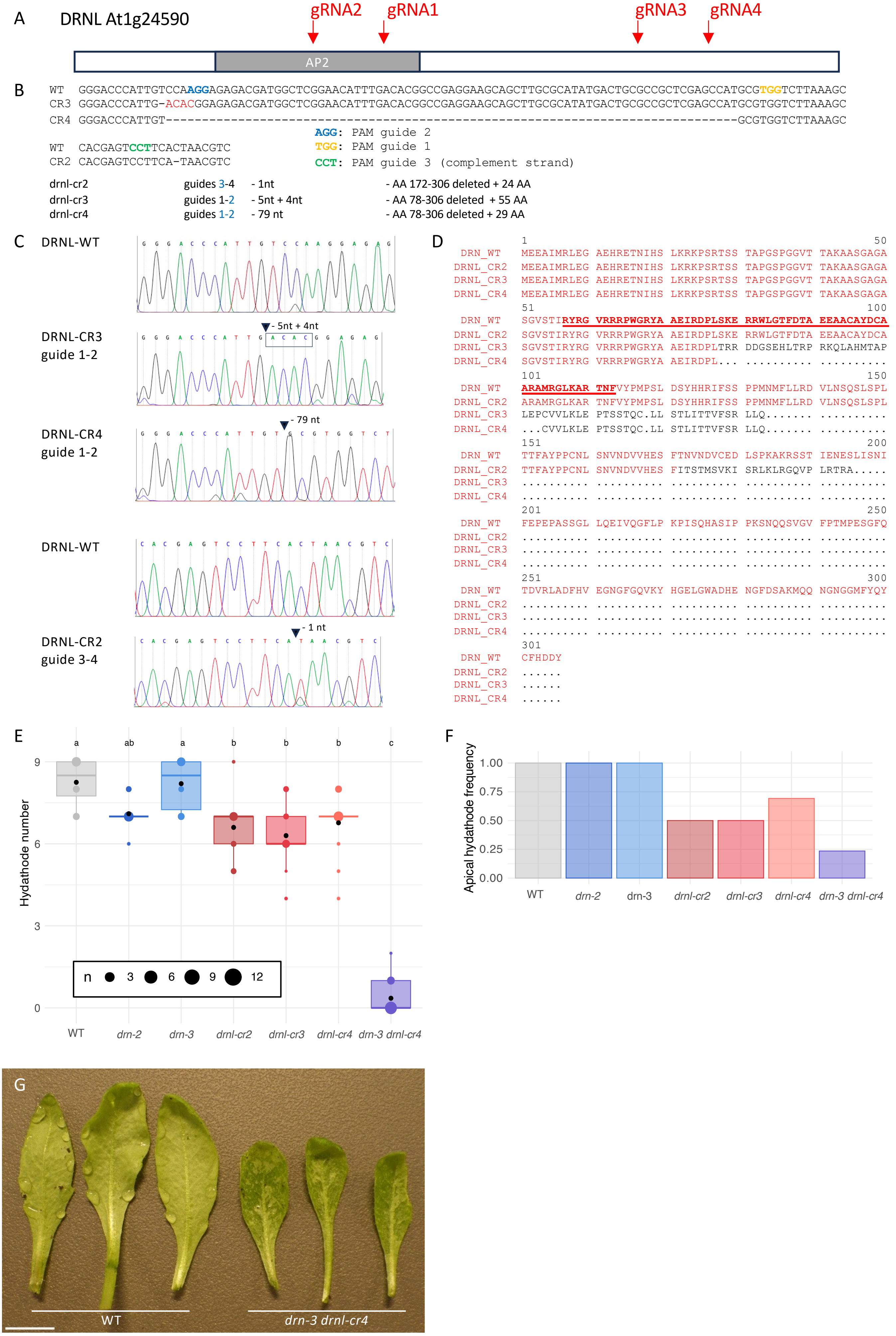
A
DRNL At1g24590
gRNA2
gRNA1
gRNA3
gRNA4
AP2
B
WT GGGACCCATTGTCCAAGGAGAGACGATGGCTCGGAACATTTGACACGGCCGAGGAAGCAGCTTGCGCATATGACTGCGCCGCTCGAGCCATGCGTGGTCTTAAAGC
CR3 GGGACCCATTG-ACACGGAGAGACGATGGCTCGGAACATTTGACACGGCCGAGGAAGCAGCTTGCGCATATGACTGCGCCGCTCGAGCCATGCGTGGTCTTAAAGC
CR4 GGGACCCATTGT---------------------------------------------------------------------------------GCGTGGTCTTAAAGC
AGG: PAM guide 2
TGG: PAM guide 1
CCT: PAM guide 3 (complement strand)
WT CACGAGTCCTTCACTAACGTC
CR2 CACGAGTCCTTCA-TAACGTC
drnl-cr2 guides 3-4 - 1nt - AA 172-306 deleted + 24 AA
drnl-cr3 guides 1-2 - 5nt + 4nt - AA 78-306 deleted + 55 AA
drnl-cr4 guides 1-2 - 79 nt - AA 78-306 deleted + 29 AA
C
DRNL-WT
DRNL-CR3 guide 1-2
- 5nt + 4nt
DRNL-CR4 guide 1-2
- 79 nt
DRNL-WT
DRNL-CR2 guide 3-4
- 1 nt
D
1 50
DRN_WT MEEAIMRLEG AEHRETNIHS LKRKPSRTSS TAPGSPGGVT TAKAASGAGA
DRNL_CR2 MEEAIMRLEG AEHRETNIHS LKRKPSRTSS TAPGSPGGVT TAKAASGAGA
DRNL_CR3 MEEAIMRLEG AEHRETNIHS LKRKPSRTSS TAPGSPGGVT TAKAASGAGA
DRNL_CR4 MEEAIMRLEG AEHRETNIHS LKRKPSRTSS TAPGSPGGVT TAKAASGAGA
51 100
DRN_WT SGVSTIRYRG VRRRPWGRYA AEIRDPLSKE RRWLGTFDTA EEAACAYDCA
DRNL_CR2 SGVSTIRYRG VRRRPWGRYA AEIRDPLSKE RRWLGTFDTA EEAACAYDCA
DRNL_CR3 SGVSTIRYRG VRRRPWGRYA AEIRDPLTRR DDGSEHLTRP RKQLAHMTAP
DRNL_CR4 SGVSTIRYRG VRRRPWGRYA AEIRDPL... .......... ..........
101 150
DRN_WT ARAMRGLKAR TNFVYPMPSL DSYHHRIFSS PPMNMFLLRD VLNSQSLSPL
DRNL_CR2 ARAMRGLKAR TNFVYPMPSL DSYHHRIFSS PPMNMFLLRD VLNSQSLSPL
DRNL_CR3 LEPCVVLKLE PTSSTQC.LL STLITTVFSR LLQ....... ..........
DRNL_CR4 ...CVVLKLE PTSSTQC.LL STLITTVFSR LLQ....... ..........
151 200
DRN_WT TTFAYPPCNL SNVNDVVHES FTNVNDVCED LSPKAKRSST IENESLISNI
DRNL_CR2 TTFAYPPCNL SNVNDVVHES FITSTMSVKI SRLKLRGQVP LRTRA.....
DRNL_CR3 .......... .......... .......... .......... ..........
DRNL_CR4 .......... .......... .......... .......... ..........
201 250
DRN_WT FEPEPASSGL LQEIVQGFLP KPISQHASIP PKSNQQSVGV FPTMPESGFQ
DRNL_CR2 .......... .......... .......... .......... ..........
DRNL_CR3 .......... .......... .......... .......... ..........
DRNL_CR4 .......... .......... .......... .......... ..........
251 300
DRN_WT TDVRLADFHV EGNGFGQVKY HGELGWADHE NGFDSAKMQQ NGNGGMFYQY
DRNL_CR2 .......... .......... .......... .......... ..........
DRNL_CR3 .......... .......... .......... .......... ..........
DRNL_CR4 .......... .......... .......... .......... ..........
301
DRN_WT CFHDDY
DRNL_CR2 ......
DRNL_CR3 ......
DRNL_CR4 ......
E
a ab a b b b c
Hydathode number
9
6
3
0
n 3 6 9 12
WT drn-2 drn-3 drnl-cr2 drnl-cr3 drnl-cr4 drn-3 drnl-cr4
F
Apical hydathode frequency
1.00
0.75
0.50
0.25
0.00
WT drn-2 drn-3 drnl-cr2 drnl-cr3 drnl-cr4 drn-3 drnl-cr4
G
WT
drn-3 drnl-cr4

**Extended Data Figure 4. Novel *drnl* alleles in Col-0 background reveal a unique role of *DRNL* in apical hydathode formation and a redundant role with *DRN* for marginal hydathode formation.**

**A.** The *DRNL* gene codes for a 306 amino-acid-long protein with a conserved AP2 domain in the N-terminal part. 4 RNA guides were designed to target *DRNL* by Cas9-CRISPR genome editing. These guides were used in couples: guide 1 + guide 2 or guide 3 and guide 4 simultaneously.

**B**. Summary of the 3 novels *drnl* mutant alleles obtained. The wild-type genomic region and the corresponding region in the mutants are shown. The PAM sequence of the 3 RNA guides leading to mutations is shown. A summary of the effects of the mutations at DNA and protein levels is indicated.

**C.** Sequencing chromatograms showing the wild-type genomic region and the corresponding region in the mutant are shown.

**D.** Predicted protein sequence produced by WT DRNL and the novel *drnl* alleles. The modified sequence is indicated in black and the AP2 domain is underlined.

**E.** Hydathode number (apical + marginal) in leaf 6 of WT, single *drn* and novel CRISPR *drnl-cr* mutants and *drn-3 drnl-cr4* double mutants ($n \geq 10$).

**F.** Frequency of apical hydathodes in leaf 6 of WT, single *drn* and novel CRISPR *drnl-cr* mutants and *drn-3 drnl-cr4* double mutants ($n \geq 10$).

**G**. Leaf flooding phenotype in *drn-3 drnl-cr4* double mutant following the induction of guttation.

Bar = 1 cm in **G**.

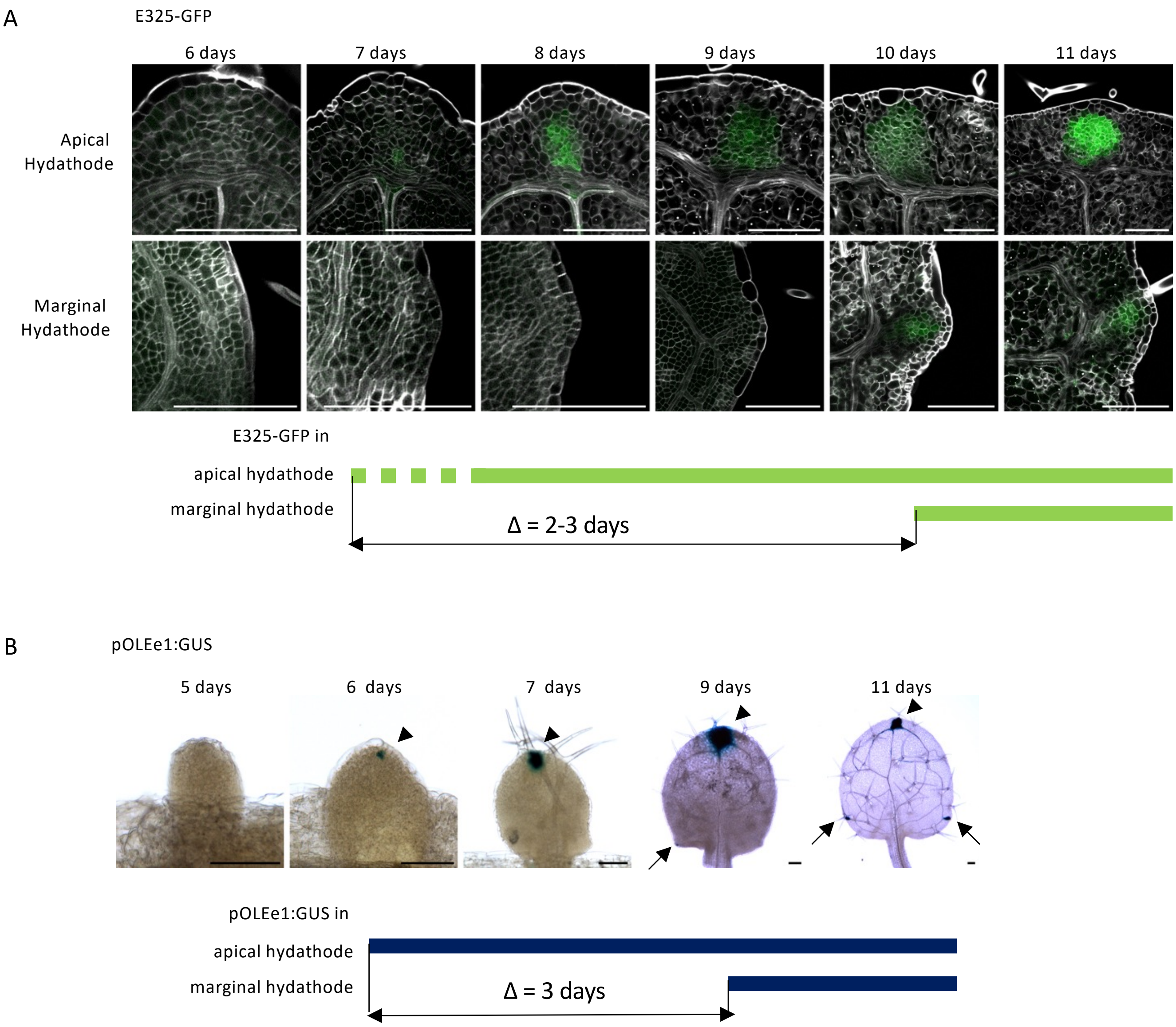


**Extended Data Figure 5. The formation of the marginal hydathodes is delayed by about 3 days compared to apical hydathodes in the first leaves.**

**A.** The expression of E325-GFP becomes visible at the apical hydathode in some samples at 7 days while at 8 days all apical hydathodes show strong E325-GFP expression. In marginal hydathodes, E325-GFP expression is visible at 10 days. In both structures, activation of E325-GFP expression occurs at the same age as differentiation of the epithem cells (the epithem cell remain smaller than the larger, expanding spongy mesophyll cells). Therefore, the timing of E325-GFP activation reveals a delay of about 2-3 days between apical and marginal hydathode formation.

**B.** The expression of pOLEe1-GFP-GUS is visible at the apical hydathode at 6 days while in marginal hydathodes pOLEe1-GFP-GUS is initiated at 9 days. Therefore, the timing of pOLEe1-GFP-GUS activation reveals a delay of about 3 days between apical and marginal hydathode formation.

GFP signal is shown in green and grey shows SR2020 cell wall staining in **A**. GUS staining appears in blue in **B**. Bars = 100 µm in all panels.

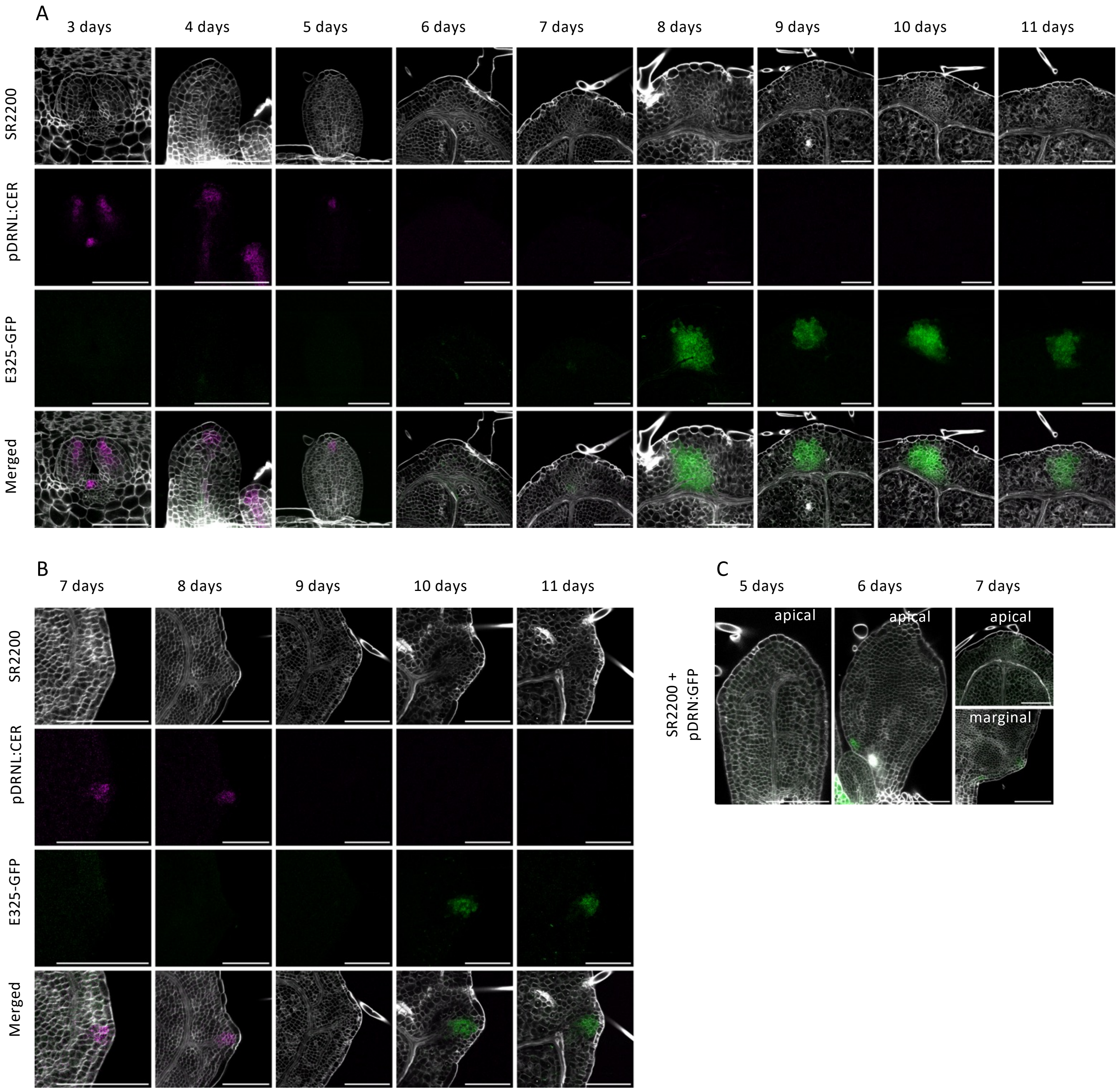


**Extended Data Figure 6. *DRNL* and *DRN* are transiently expressed at the incipient hydathode sites.**

**A.** In the leaf apex, pDRNL:CER is expressed at the incipient apical hydathode sites from day 3 to day 5, before any visible cellular differentiation of the epithem cells or activation of the E325-GFP reporter.

**B**. At the leaf margin, pDRNL:CER is expressed at the incipient marginal hydathode sites at day 7 and day 8 before any visible cellular differentiation of the epithem cells or activation of the E325-GFP reporter.

**C.** No expression of the pDRN:GFP reporter was observed at the leaf apex, while it was transiently expressed at the incipient marginal hydathode sites at day 7 and day 8.

CER signal is shown in magenta in **A** and **B**, GFP signal is shown in green in **A**-**C** and grey shows SR2020 cell wall staining in **A**-**C**. Bars= 100 µm in all panels.

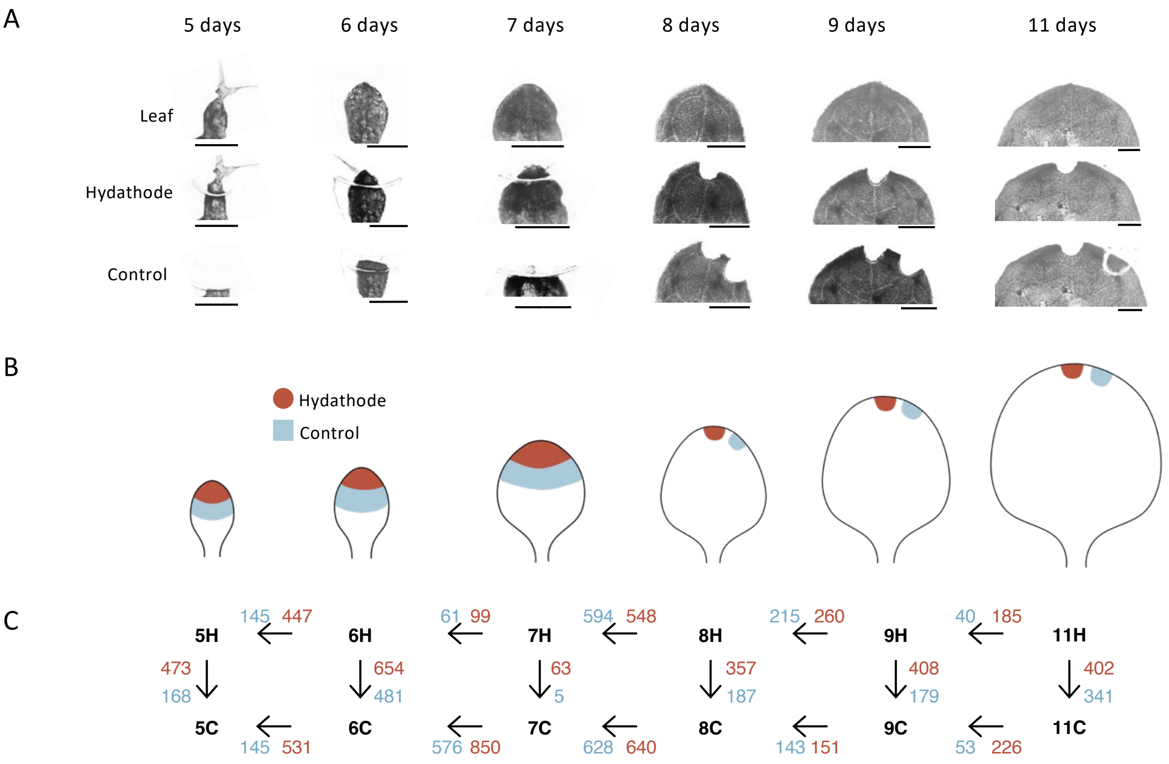


D

| Gene ID | Description TAIR | 5 days | | 7 days | | 9 days | | 11 days | |
|---|---|---|---|---|---|---|---|---|---|
| | | T | R | T | R | T | R | T | R |
| At3g51060 | SHI RELATED SEQUENCE 1; SRS1; STY1; STYLISH 1 | | | | | | | | |
| At4g36260 | SHI RELATED SEQUENCE 2; SRS2; STY2; STYLISH 2 | | | | | | | | |
| At3g16670 | Pollen Ole e 1 allergen and extensin family protein | | | | | | | | |
| At3g05730 | DEFL205; defensin-like protein 205 | | | | | | | | |
| At2g38940 | ARABIDOPSIS THALIANA PHOSPHATE TRANSPORTER 2; ATPT2; PHOSPHATE TRANSPORTER 1;4; PHT1;4 | | | | | | | | |
| At1g56710 | PGL1;POLYGALACTURONASE LIKE 1, | | | | | | | | |
| At1g08090 | ACH1; ATNRT2.1; ATNRT2:1; LATERAL ROOT INITIATION 1; LIN1; NITRATE TRANSPORTER 2 | | | | | | | | |
| At3g28740 | CYP81D11; cytochrome P450, family 81, subfamily D, polypeptide 11 | | | | | | | | |

Expressed in hydathodes based on transcriptomic data
Expressed in hydathodes based on reporter line

E

| | 5 days | 6 days | 7 days | 8 days | 9 days | 11 days |
|---|---|---|---|---|---|---|
| *STY1* | | | | | | |
| *STY2* | | | | | | |
| *SHI* | *filter* | *filter* | ns | | | ns |
| *LRP1* | ns | ns | *filter* | *filter* | *filter* | *filter* |
| *SRS5* | ns | | *filter* | ns | ns | |
| *SRS7* | ns | | *filter* | *filter* | ns | |
| *YUC1* | *filter* | *filter* | *filter* | NoObs | NoObs | NoObs |
| *YUC2* | ns | | ns | ns | ns | ns |
| *YUC4* | ns | ns | ns | ns | *filter* | *filter* |

Gene overexpressed in hydathode

F

| | 5 days | 6 days | 7 days | 8 days | 9 days | 11 days |
|---|---|---|---|---|---|---|
| *PGL1* | *filter* | *filter* | ns | | | |
| *EXLA2* | ns | | ns | | ns | |

Gene overexpressed in hydathode

Extended Data Figure 7

**Extended Data Figure 7. Transcriptomic profiling of Laser-Assisted Microdissected developing hydathodes.**

**A.** Representative hydathode and control samples collected using Laser-Assisted Microdissection in the first pair of developing leaves in 5-, 6-, 7-, 8-, 9- or 11-day-old seedlings. Control tissues are collected below the hydathode sample at 5, 6, and 7 days, and at the margin at 8, 9 and 11 days. Scale bars = 150 µm at 5 or 6 days, 300 µm at 7 days and older samples.

**B.** Scheme of the hydathode and control samples collected at different time points.

**C.** Number of genes more expressed (in red) or less expressed (in blue) for each comparison indicated by an arrow.

**D**. The expression dynamics of eight genes in developing hydathodes are similar in transcriptomic data and in reporter line analyses.

**E**. Expression of six members of the *SHI*/*STY* gene family in the transcriptomic data. *STY1* and *STY2* are overexpressed in the hydathode at all time points in the time course. The others are either overexpressed only at specific time points or not differentially expressed.

**F**. Expression of *PGL1* and *EXLA2* in the transcriptomic data.

In **E** and **F**, not differentially expressed (ns), filtered out (filter) or not detected (NoObs).

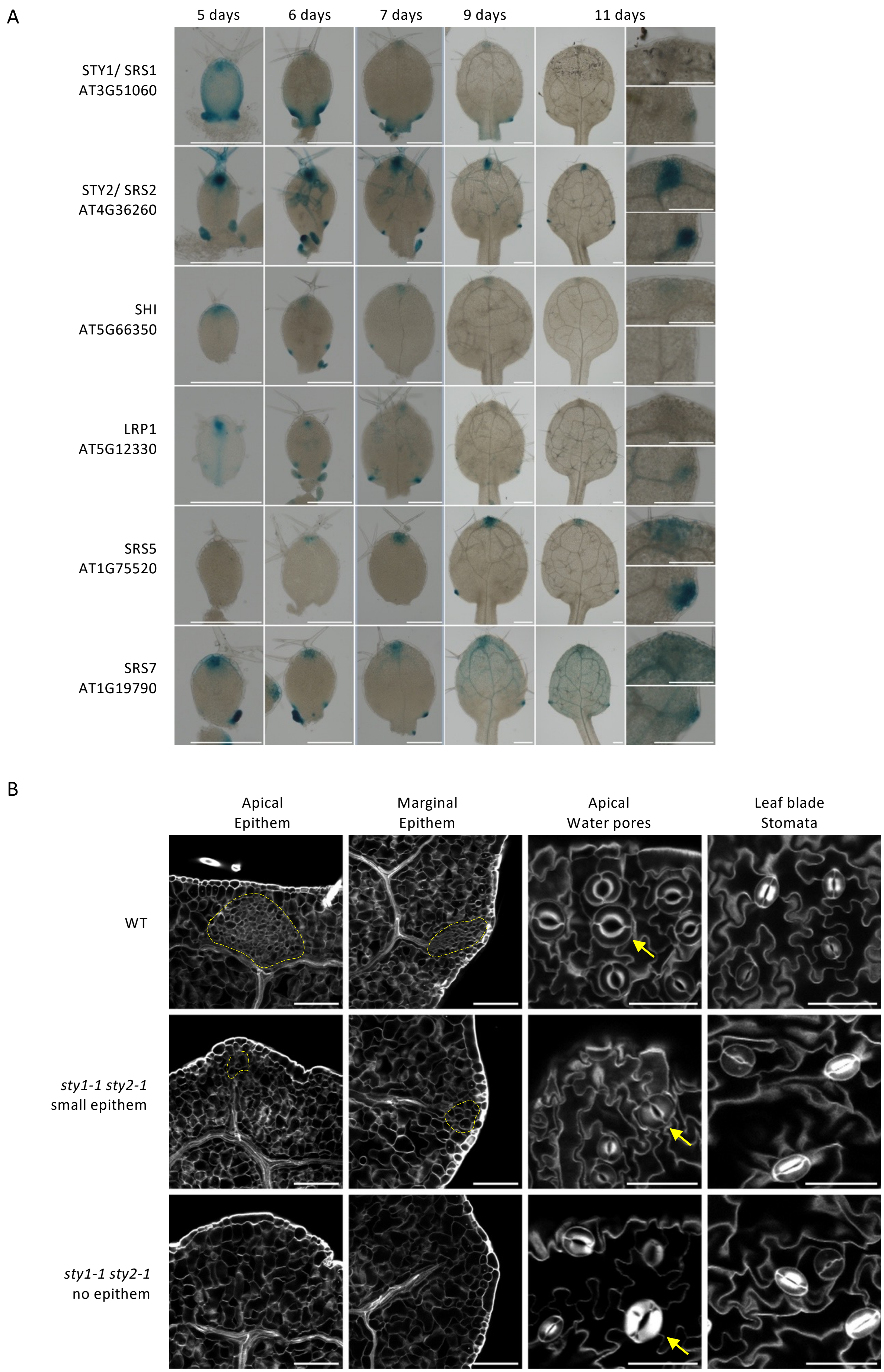

A
5 days
6 days
7 days
9 days
11 days
STY1/ SRS1
AT3G51060
STY2/ SRS2
AT4G36260
SHI
AT5G66350
LRP1
AT5G12330
SRS5
AT1G75520
SRS7
AT1G19790
B
Apical
Epithem
Marginal
Epithem
Apical
Water pores
Leaf blade
Stomata
WT
sty1-1 sty2-1
small epithem
sty1-1 sty2-1
no epithem

**Extended Data Figure 8. *STY1* and *STY2* are required for hydathode formation.**

**A.** Expression patterns of 6 members of the *SHORT INTERNODES/STYLISH* gene family during hydathode formation in leaf 1. All the genes analyzed are expressed in developing apical and marginal hydathodes, although with different kinetics. For instance, *STY1* and *STY2* are both expressed early on, but *STY2* expression remains strong until day 11, while *STY1* expression decreases. On the opposite, *SRS5* expression starts later (at day 6 in the apical hydathode), but remains high until day 11.

**B.** The *sty1-1 sty2-1* double mutant shows hydathodes with smaller epithem than the WT at both apical and marginal positions. In some leaves, no sign of differentiated epithem is observed. A few water pores can be observed at the *sty1-1 sty2-1* leaf apex.

In **A**, the last column shows a detail of the apical and marginal hydathodes of day 11 leaves. In **B**, grey shows SR2200 cell wall staining, the epithem is circled by a yellow dotted line and yellow arrows point to water pores. Bars = 200 µm in **A**, and 100 µm in **B.**

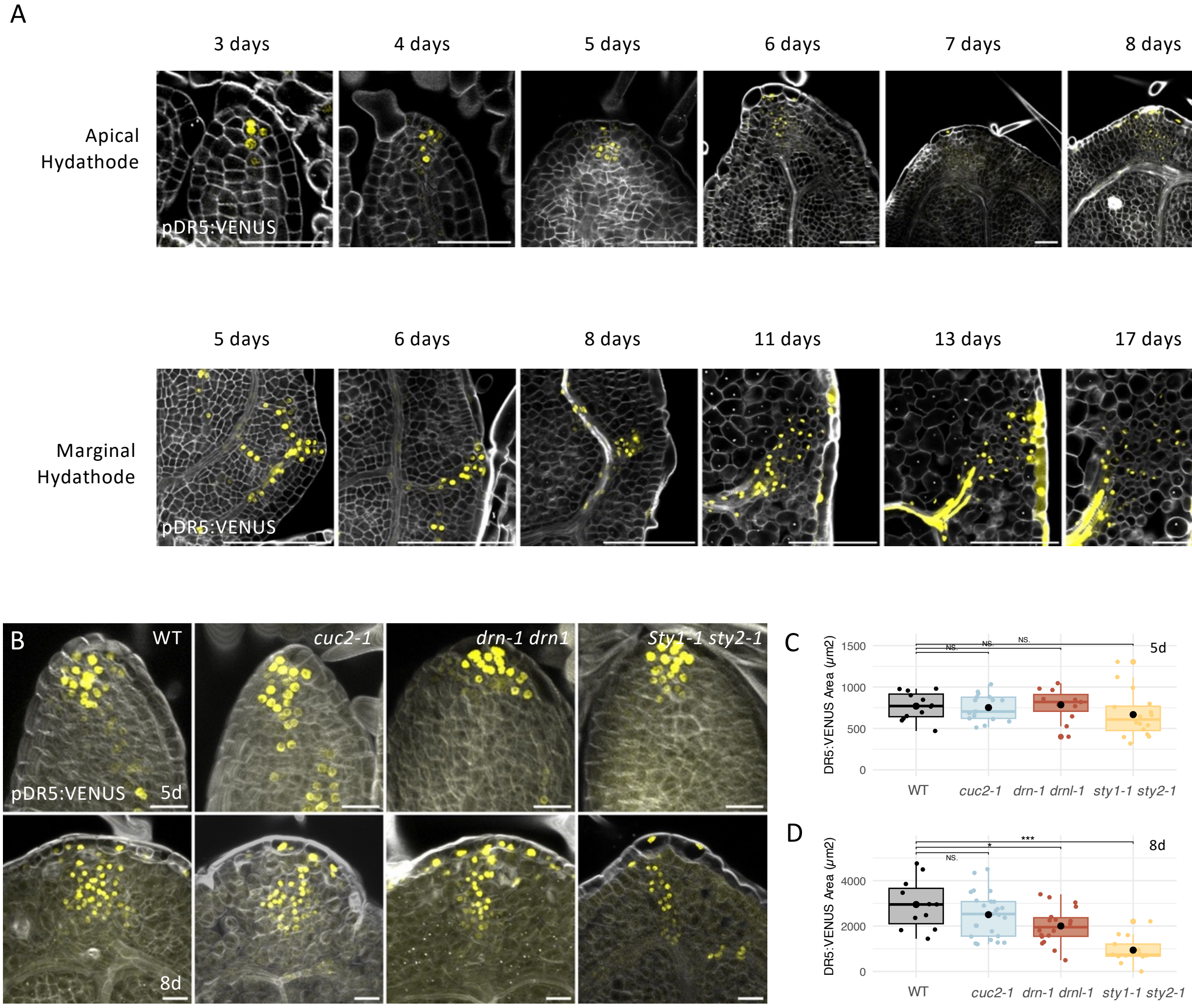


**Extended Data Figure 9. Auxin signaling occurs throughout hydathode formation.**

**A.** pDR5:VENUS expression during apical and marginal hydathode formation. Auxin signaling is observed in hydathodes from their initiation stage through to maturity.

**B.** pDR5:VENUS expression at the apical hydathode site at 5 days (upper panels) or 8 days (lower panels), in WT, *cuc2-1* mutants or *drn-1 drnl-1* and *sty1-1 sty2-1* double mutants.

**C.** The area of pDR5:VENUS expression domain is unchanged in the *cuc2-1* mutant and *drn-1 drnl-1* and *sty1-1 sty2-1* double mutants during the specification of apical hydathodes (5 days) ($n \geq 11$).

**D.** The area of pDR5:VENUS expression domain is unchanged in the *cuc2-1* but reduced in *drn-1 drnl-1* and *sty1-1 sty2-1* double mutants during the differentiation of apical hydathodes (8 days) ($n \geq 11$).

VENUS signal is shown in yellow and SR2200 cell wall staining is shown in grey in all panels. Bars = 50 µm in **A** Apical and 100 µm in **A** Marginal, 20 µm in **B**.

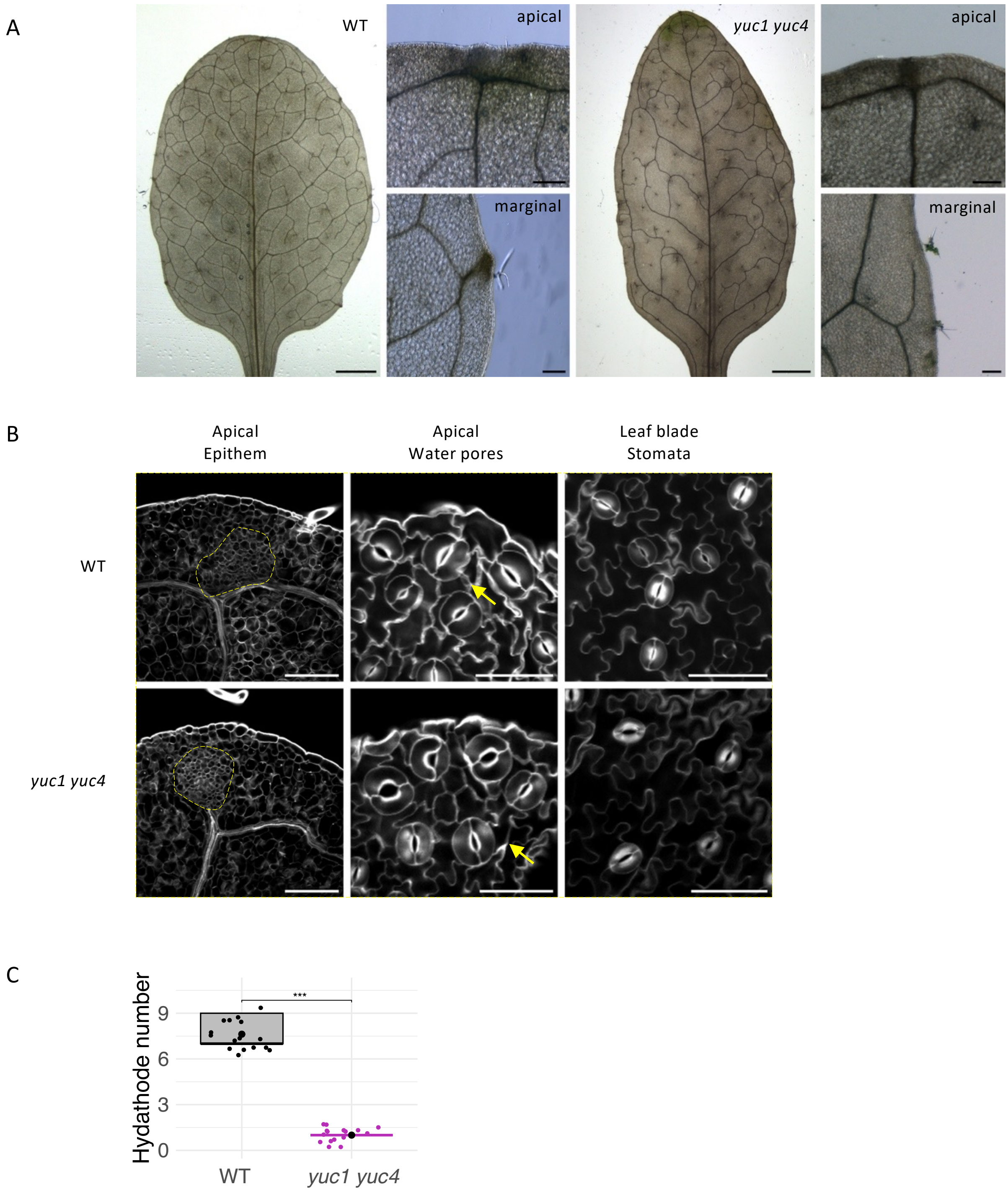


**Extended Data Figure 10. *YUC1* and *YUC4* are required for hydathode formation.**

**A.** Ethanol-cleared leaf 6 of the WT and *yuc1 yuc4* double mutant. The *yuc1 yuc4* double mutant develops an apical hydathode but lacks marginal hydathodes**.**

**B**. A clear epithem is visible at the apical hydathode and several water pores are present in the *yuc1 yuc4* double mutant.

**C.** The number of hydathodes is reduced in *yuc1 yuc4* leaf 6 compared to the WT ($n$ = 16).

In **B**, grey shows SR2200 cell wall staining, the epithem in circled by a yellow dotted line and yellow arrows point to water pores. Bars = 2 mm in **A**, and 100 µm in **B.**

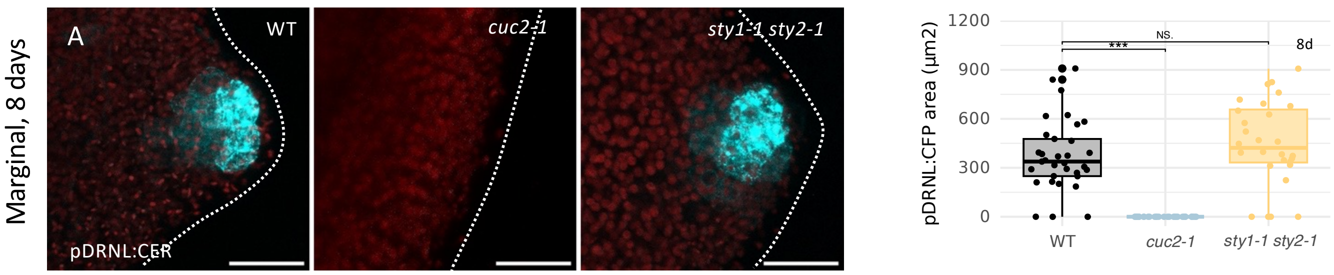


**Extended Data Figure 11. *DRNL* expression requires *CUC2* but is independent of *STY1/STY2* at incipient marginal hydathode sites.**

**A.** Expression of pDRNL:CER at the incipient marginal hydathode site at 8 days in WT, *cuc2-1* mutant and *sty1-1 sty2-1* double mutants.

**B.** The area of pDRNL:CER expression domain is unchanged in the *sty1-1 sty2-1* double mutant during the specification of marginal hydathodes (8 days) (*n* = 28). No expression of pDRNL:CER is observed at the leaf margin of *cuc2-1* mutants at 8 days.

In **A**, CER signal is shown in cyan and chlorophyl fluorescence is shown in red. Bars = 20 µm in all panels**.**

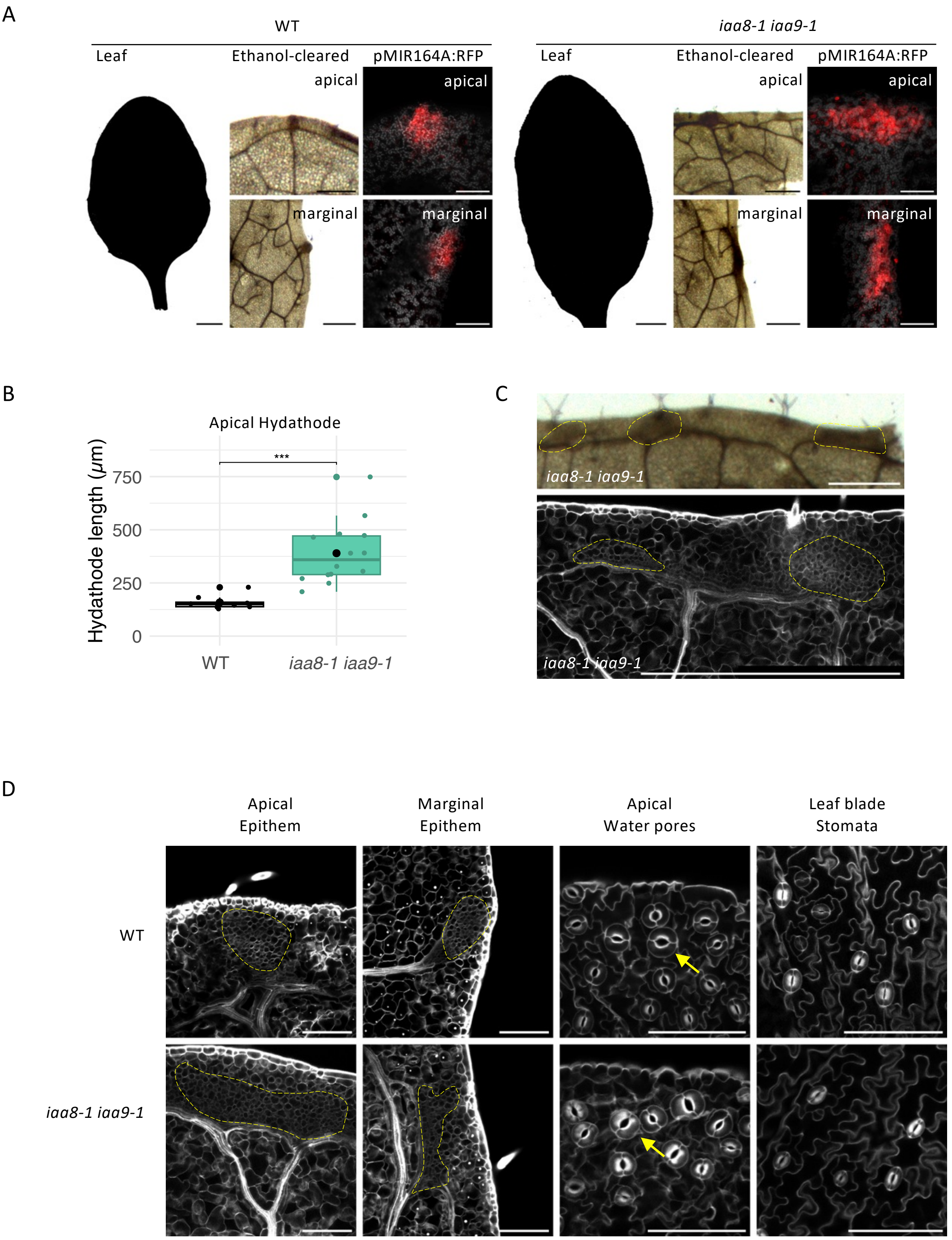


**Extended Data Figure 12. Extra and larger hydathodes form in *iaa8 iaa9* double mutants.**

**A.** Leaf shape and hydathodes in WT and *iaa8-1 iaa9-1* double mutants. For each genotype, a mature leaf 6 silhouette is shown in the first panel, the second and third panels show ethanol-cleared apical and marginal hydathodes, and the fourth and last panel show pMIR164A:RFP expression as a marker of hydathodes in apical and marginal hydathodes. The *iaa8-1 iaa9-1* double mutants have smooth leaves with larger expression domains of pMIR164A:RFP in both apical and marginal hydathodes.

**B.** Leaf 6 apical hydathode are longer in *iaa8-1 iaa9-1* compared to WT, using pMIR164A:RFP expression domain as a proxy for hydathodes ($n \geq 9$).

**C.** Ectopic hydathodes develop in the apical region of *iaa8-1 iaa9-1*. The upper panel shows an ethanol-cleared leaf with 3 hydathodes developing in the apical region. The lower panel shows with a cell wall-stained leaf with 2 hydathodes developing in the apical region

**D.** Larger apical and marginal hydathodes are observed in *iaa8-1 iaa9-1* compared to WT. Water pores are located above the apical hydathode in *iaa8-1 iaa9-1.*

In **C** and **D**, grey shows SR2200 cell wall staining, the epithem in circled by a yellow dotted line and yellow arrows point to water pores. Bars = 2 mm for the entire leaves in **A**, 0.5 mm for the ethanol-cleared panels in **A** and both panels in **C**, and 100 μm for the confocal panels in **A** and **D**.

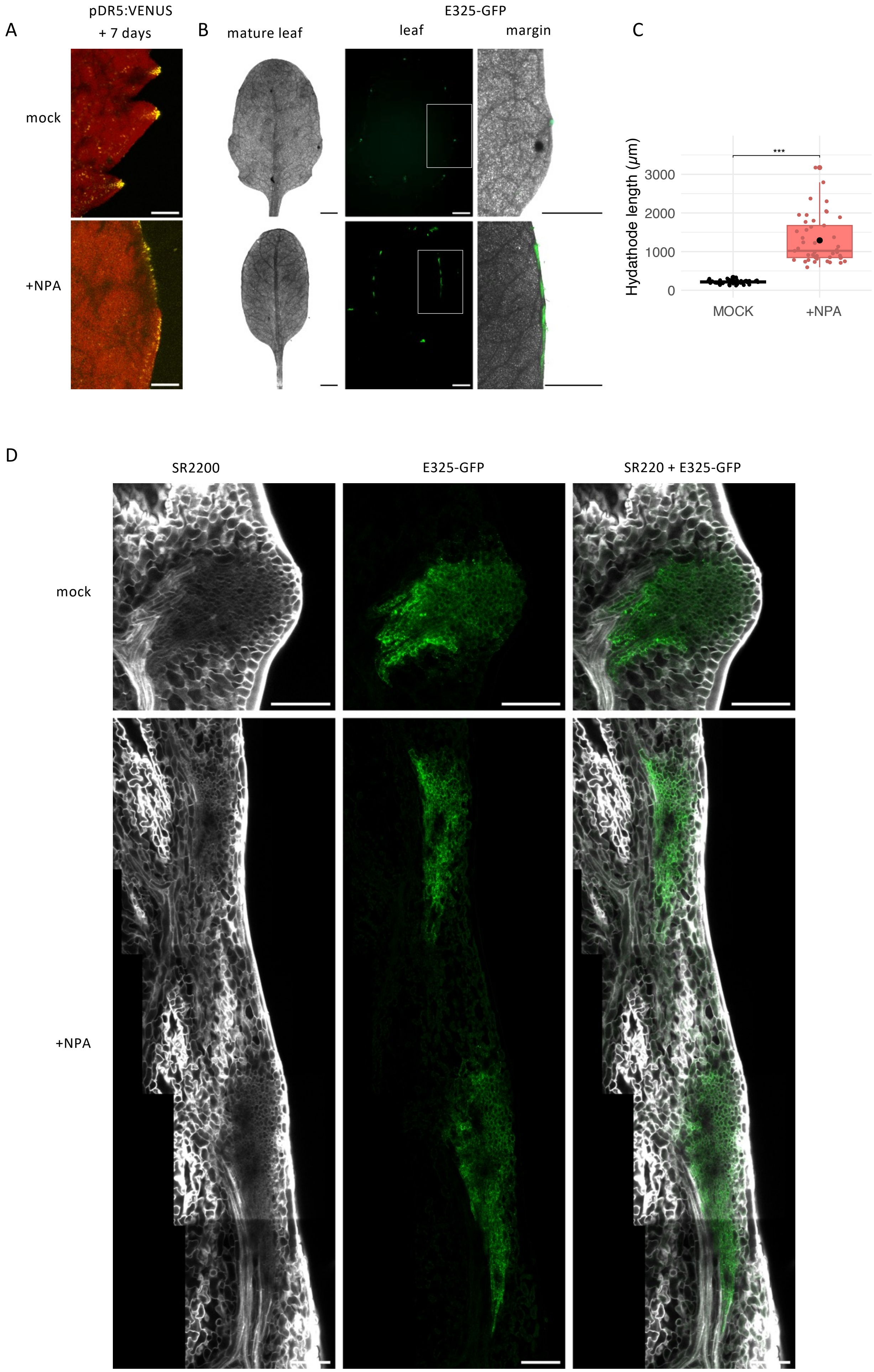


Extended Data Figure 13

Savourat et al.

**Extended Data Figure 13. NPA-treated leaves form large hydathodes.**

**A.** Detail of leaf margins showing pDR5:VENUS expression as a proxy of auxin response. Leaves were observed 7 days following NPA application. In the mock-treated leaves, localized DR5-VENUS expression is observed while in NPA-treated plants pDR5:VENUS expression appears more continuous along the margin.

**B.** E325-GPP expression as a hydathode maker in mature mock- or NPA-treated leaves. Following NPA application, leaves develop smoothed margins in contrast to the serrated mock-treated leaves (first panel of each row). In mock-treated leaves, E325-GFP forms small dots of expression, while in NPA-treated leaves, E325-GFP forms elongated domains along the leaf margin (second and third panels of each row).

**C.** In NPA-treated leaves, marginal hydathodes are longer than in mock-treated leaves, using E325-GFP expression as a proxy for hydathodes ($n \geq 45$).

**D.** Detail of hydathode organization in mature mock- or NPA-treated leaves. Enlarged epithems marked by E325-GFP and connected to the vasculature network develop in NPA-treated leaves.

pDR5:VENUS signal is shown in yellow and chlorophyll fluorescence in red in **A**. E325-GFP is shown in green in **B** and **C**, chlorophyll fluorescence in grey in **B** and SR2200 cell wall staining in grey in **D**. Bars = 100 µm in **A**, 2 mm in **B**, 100 µm in **D**.

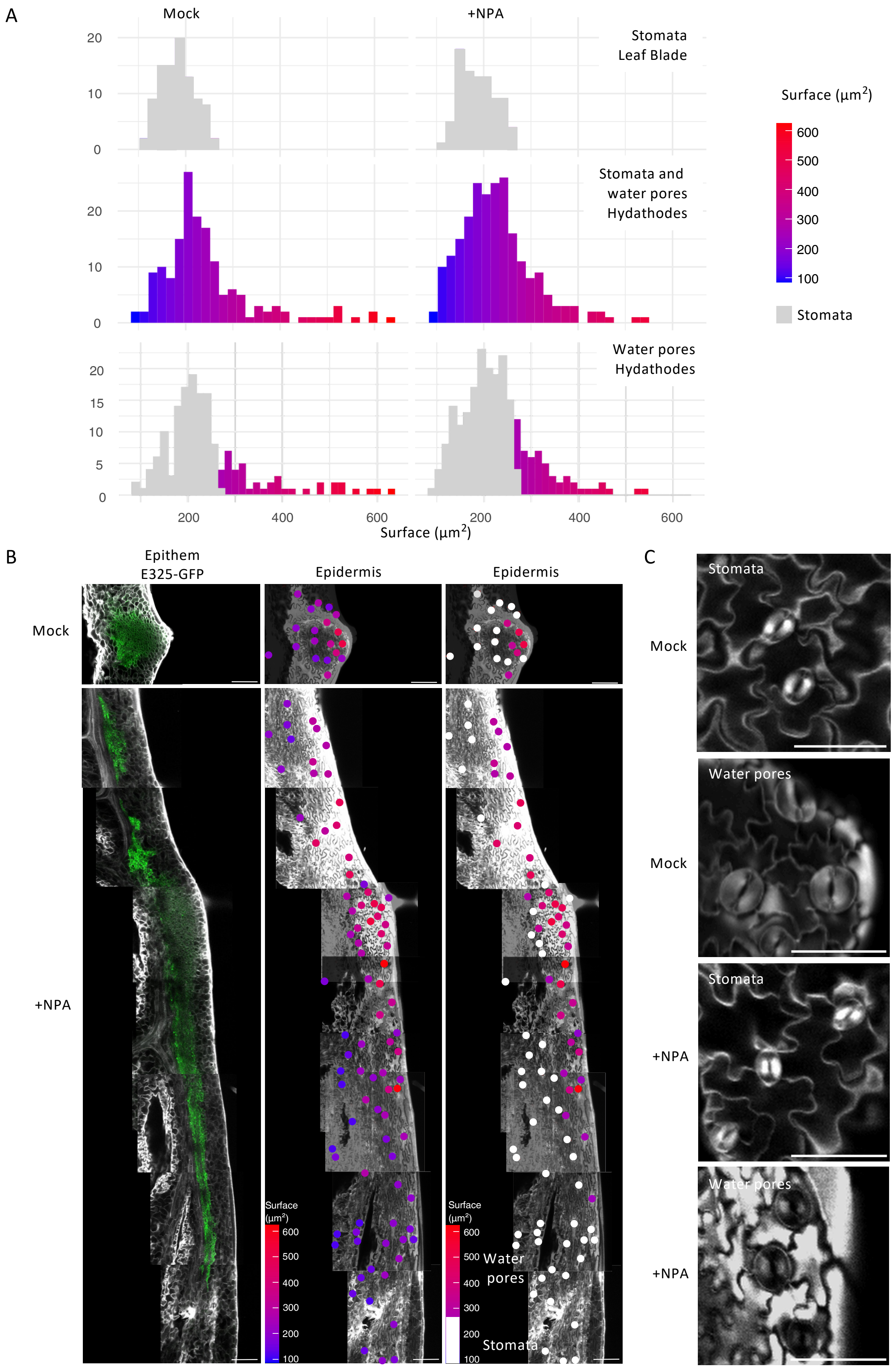
A
Mock
+NPA
Stomata
Leaf Blade
Stomata and
water pores
Hydathodes
Water pores
Hydathodes
Surface (µm²)
600
500
400
300
200
100
Stomata
Surface (µm²)
B
Epithem
E325-GFP
Epidermis
Epidermis
Mock
+NPA
Water
pores
Stomata
C
Stomata
Water pores
Mock
+NPA

**Extended Data Figure 14. The enlarged hydathodes of NPA-treated leaves are covered by water pores.**

**A.** Strategy to identify water pores. The size of true stomata was measured at the blade center in mock- and NPA-treated leaves (first line). Stomata and putative water pores size was measured in the hydathode region in mock- and NPA-treated leaves (second line). Water pores were determined as structures located close to hydathodes and which size is bigger than the one of true stomata (third line).

**B.** The panels in the first column show an optical section through the epithem expressing the E325-GFP reporter in a mock and NPA-treated leaf. In the panels of the second column showing the epidermis, stomata-water pores are colored according to their size. The panels of the last column show size-based color coding of water pores and stomata (in grey). Note that water pores are located in an elongated region along the epithem in the NPA treated leaf.

**C.** Detail of representative stomata and water pores of mock- and NPA-treated leaves.

E325-GFP is shown in green in **B**, SR2200 cell wall staining in grey in **B** and **C**. Bars = 2 mm in **B**, 50 µm in **C**.

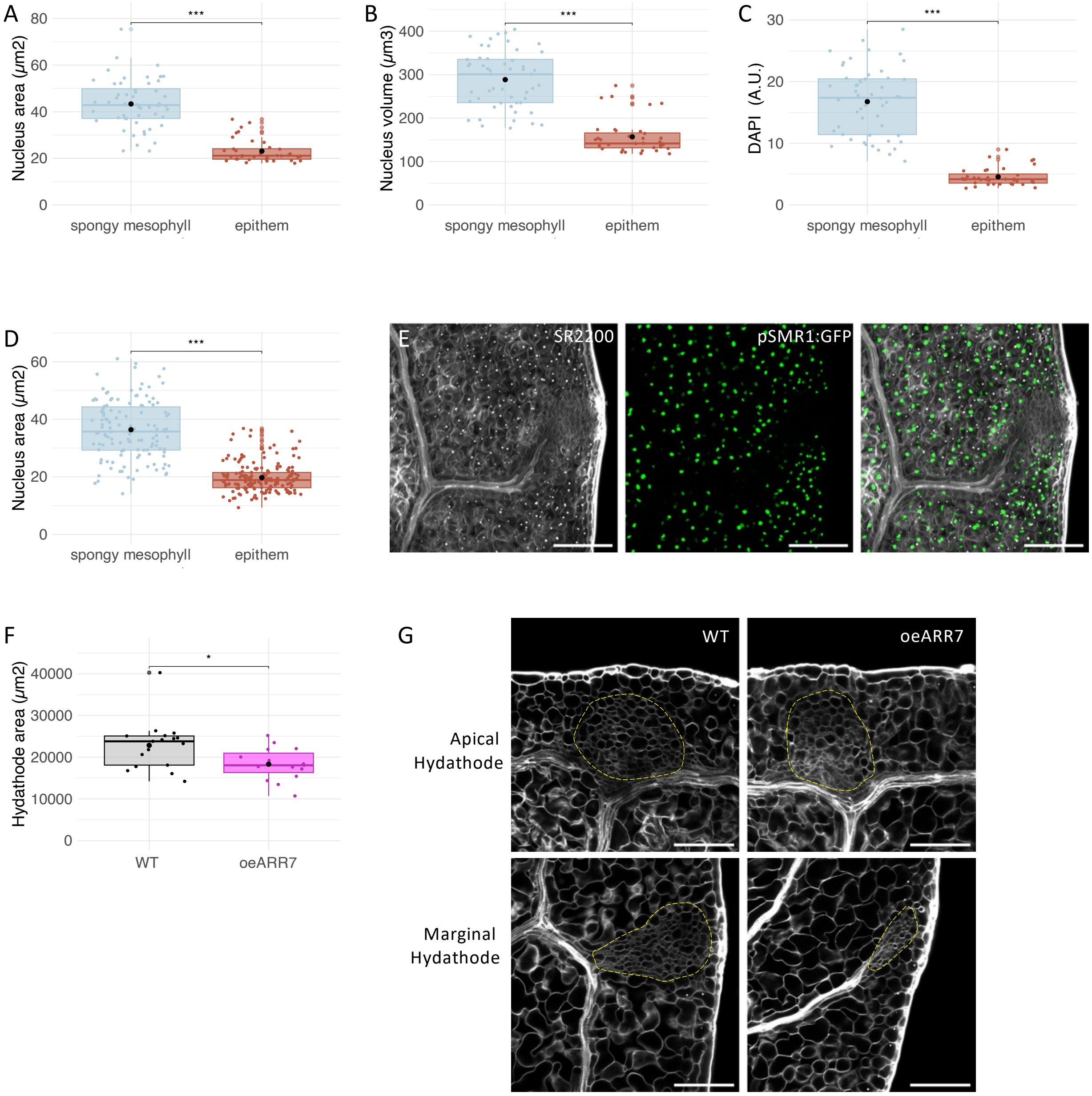


**Extended Data Figure 15. Endoreduplication is reduced in epithem cells.**

**A.** In apical hydathodes of leaf 1 in 17-day-old seedlings, the nucleus area is smaller in epithem cells than in spongy mesophyll cells (*n* ≥ 102).

**B**. In apical hydathodes of leaf 1 in 17-day-old seedlings, the nucleus volume is smaller in epithem cells than in spongy mesophyll cells (*n* ≥ 36).

**C**. In apical hydathodes of leaf 1 in 17-day-old seedlings, the DAPI signal is lower in epithem cells than in spongy mesophyll cells (*n* ≥ 36).

**D.** In marginal hydathodes of leaf 1 in 17-day-old seedlings, the nucleus area is lower in epithem cells than in spongy mesophyll cells (*n* ≥ 126).

**E.** The endoreduplication marker pSMR1:GFP is not expressed in the marginal hydathode region of leaf 1 at 17 days.

**F.** In leaf 1 of 17-day-old seedlings, the apical hydathode size is smaller in a ARR7 overexpressing line (oeARR7) compared to WT, using epithem size measured on SR2200-stained sections as a proxy for hydathode size (*n* ≥ 15).

**G.** Hydathodes of a ARR7 overexpressing line (oeARR7) show smaller apical and marginal hydathodes.

SR2200 cell wall staining is shown in grey in **E** and **G** and GFP signal is shown in green in **E**. The epithem in circled by a yellow dotted line in **G**. Bars = 100 µm in **E** and **G**.

A

B

C

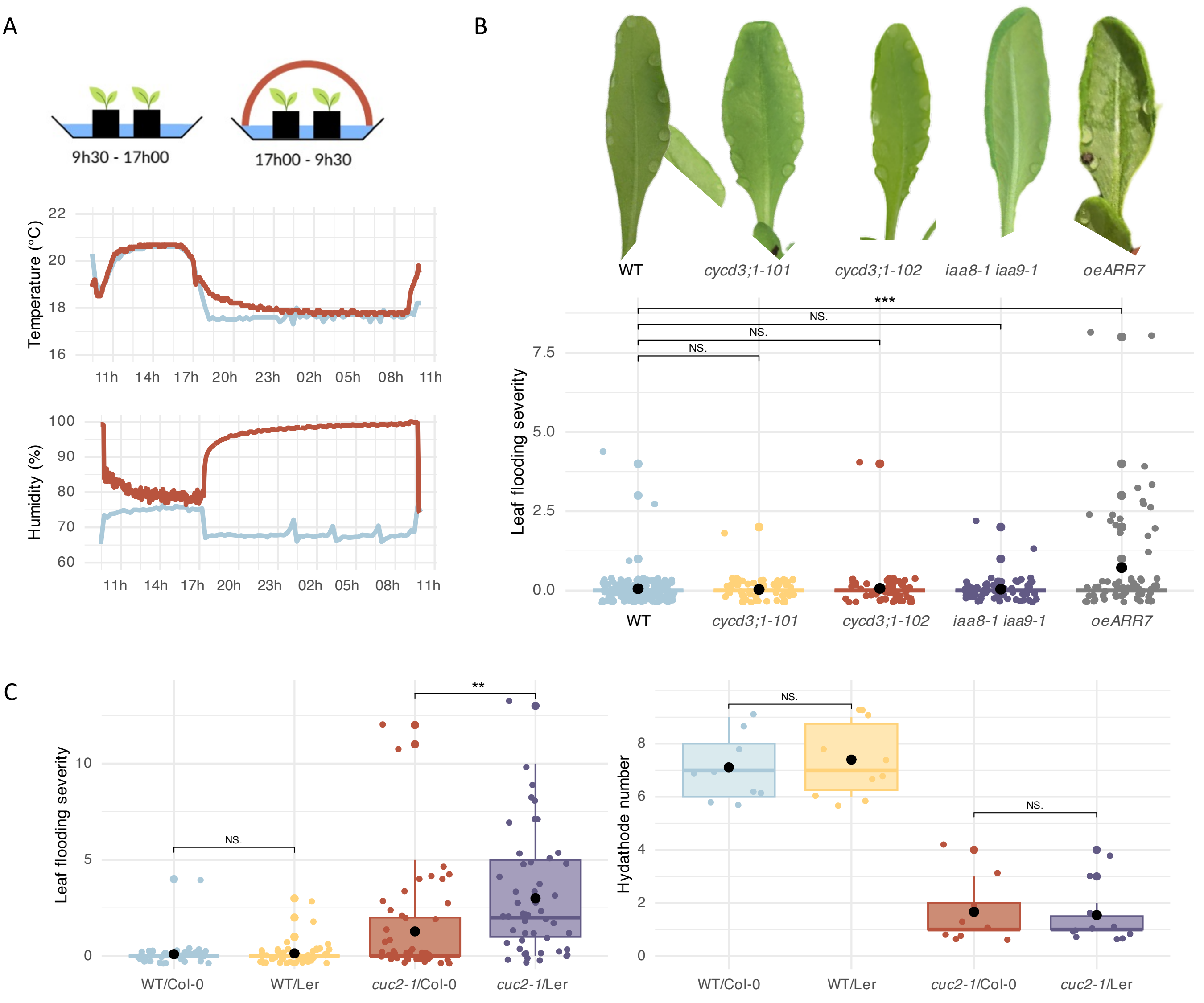


D

| | Week | Leaf Rank | | | | | | | | | | | |
|---|---|---|---|---|---|---|---|---|---|---|---|---|---|
| | | 1 | 2 | 3 | 4 | 5 | 6 | 7 | 8 | 9 | 10 | 11 | 12 |
| Leaf flooding severity | 3 | 0.50 | 0.50 | 0.00 | 0.00 | 0.00 | 0.00 | 0.00 | NA | NA | NA | NA | NA |
| | 4 | 2.88 | 2.88 | 6.25 | 3.00 | 0.50 | 0.00 | 0.00 | 0.00 | 0.00 | 0.00 | 0.00 | 0.00 |
| | 5 | 3.63 | 4.38 | 5.00 | 5.25 | 4.00 | 6.50 | 3.75 | 1.25 | 2.75 | 2.00 | 0.25 | 0.00 |
| | 6 | 0.50 | 0.50 | 1.25 | 6.75 | 5.75 | 2.25 | 2.75 | 3.00 | 4.00 | 3.25 | 3.25 | 2.25 |

0 1 2 3 4 5 6 7 8 9 10 11 12 13 14 15 16 17 18 19 20

E

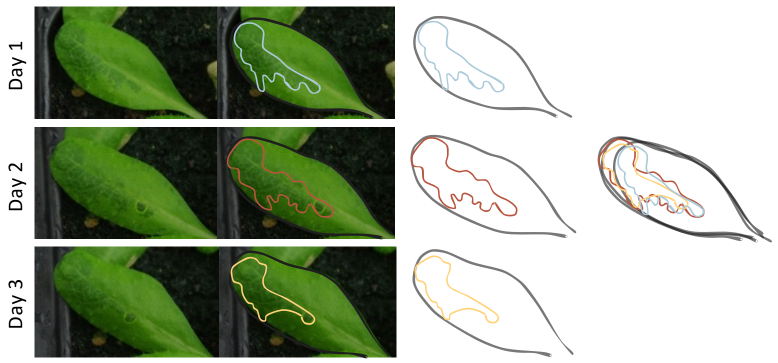

**Extended Data Figure 16. Characterization of leaf flooding in hydathode mutants.**

**A**. Protocol to induce leaf guttation/flooding. Well-watered plants are covered by a lid over-night and guttation/flooding is observed in the next morning immediately after lid removal.

**B**. Leaves following over-night induction of guttation show guttation droplets or leaf flooding (visible as dark patches on the leaf blade). Leaves of WT, *cycd3;1-101* and *cycd3;1-102* mutants, *iaa8-1 iaa9-1* double mutants and *ARR7* overexpressors are shown. The quantification of leaf flooding severity in these lines shows that oeARR7 has weakly flooded leaves. Leaf flooding severity is indicated by a score ranging from 0 (no flooding) to 20 (100% of the leaf blade area flooded) ($n \geq 59$).

**C**. Quantification of leaf flooding severity in WT/Col-0, WT/L*er cuc2*-1/Col-0 and *cuc2-1*/L*er*. When the same *cuc2-1* allele is present in the L*er* background, it leads to more severe leaf flooding than when present in Col-0 background (left panel, $n \geq 40$). Note that the number of hydathodes in leaf 6 is similar in both genetic backgrounds (right panel, $n \geq 9$).

**D.** Quantification of flooding severity in *cuc2-1*/L*er* leaves of different ranks and plants with different ages (from 3- to 6-week-old). For a given leaf, severity of flooding changes during its maturation. For instance, leaves 1 and 2 show increasing leaf flooding severity from 3- to 5- weeks followed by a strong reduction ($n = 4$).

**E.** Flooding repeats along similar patterns if induced over several days. Guttation was induced for 3 consecutive days in *cuc2-1*/L*er* leaves. Blue: flooded area on day 1, red: flooded area on day 2, and yellow: flooded area on day 3. Gray outline: leaf shape.

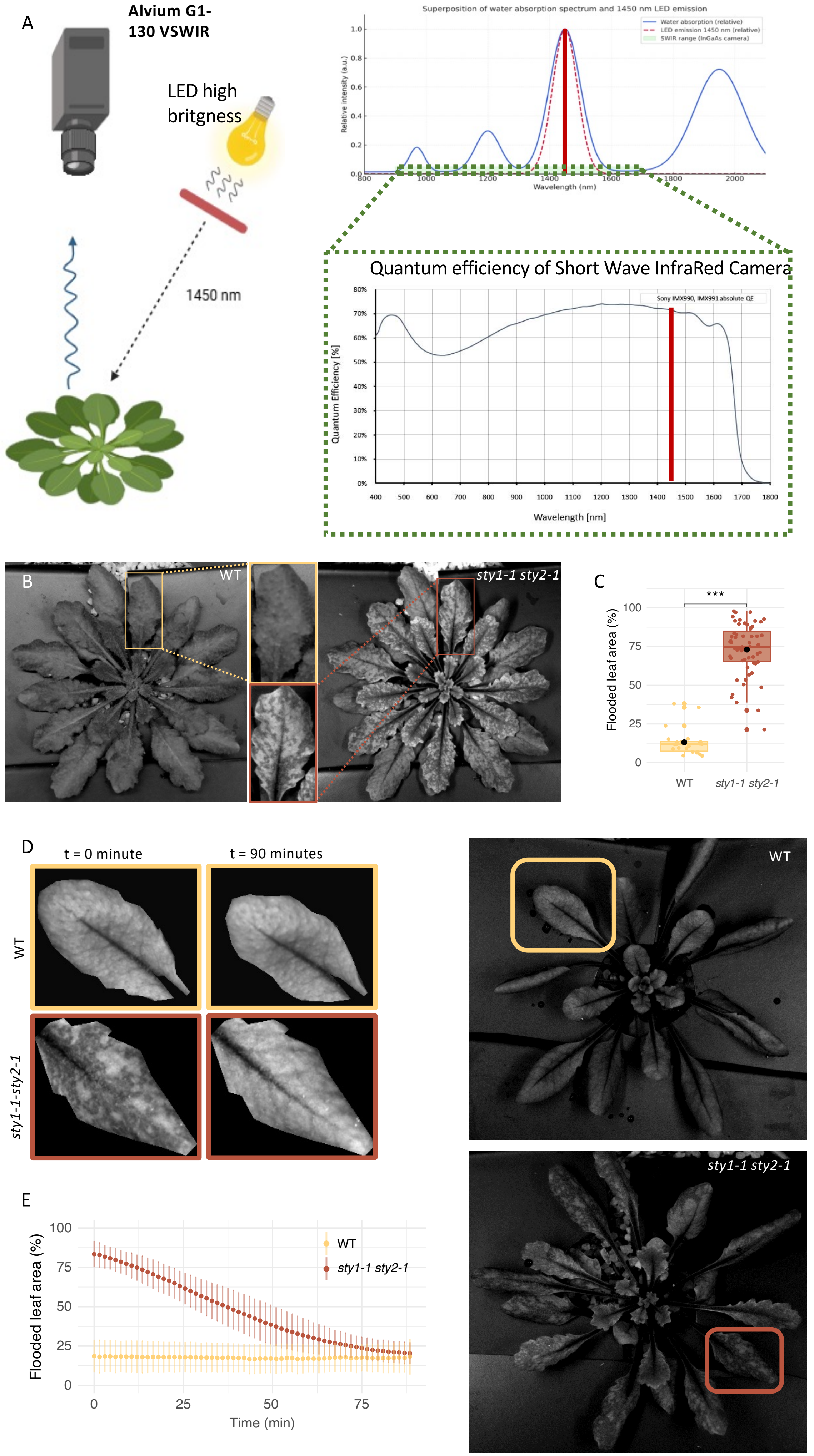


Extended Data Figure 17

Savourat et al.

**Extended Data Figure 17. Flooding imaging using a Short Wave InfraRed Camera (SWIR).**

**A.** SWIR Imaging experimental set-up. Hight brightness LED light up with a bandwidth at 1450 nm. At this wavelength, flooded tissues absorb all the light because of water absorption spectrum. Dry tissues reflected LED light at 1450 nm which is visible to the SWIR camera. Dry tissues correspond to clear area and flooded tissues correspond to dark area on the leaf.

Superposition of water absorption spectrum (blue line) and LED EFFI-FLEX-5-1450 emission (red dotted line). Maximal water absorption is around 1450 nm. In green box below, quantum efficiency of G-130-VSWIR which absorbs between 400 nm and 1700 nm.

**B.** WT and *sty1-1 sty2-1* double mutant rosettes following induction of flooding/guttation and imaged using the SWIR system. Flooded areas appear as dark patches in the *sty1-1 sty2-1* double mutant.

**C.** Quantification of the flooded area in WT and *sty1-1 sty2-1* double mutant (expressed as percentage of total leaf area) ($n \geq 22$).

**D.** Flooding reversion in *sty1-1 sty2-1* double mutant and for comparison WT. Rosettes were removed from the flooding/guttation inductive conditions and imaged for 90 minutes using the SWIR device. Details show a rosette leaf of WT or *sty1-1 sty2-1* double mutant at t=0 or t=90 min, while the full rosettes at t=0 is shown on the right. Movies are available as Extended Data Movie 1 and 2.

**E.** Kinetic of flooding reversion. Flooding is shown as the percentage of total leaf area, bars represent the standard deviation ($n \geq 10$). Note, that at the end of the *sty1-1 sty2-1* reversion kinetic as well as in the WT, the value remains about 20% which is due to the presence of the vasculature that has a high content in water and appears as dark regions.

| Reporter/Dye | Excitation wavelength (nm) | Detection wavelength (nm) |
|---|---|---|
| pMIR164A:erRFP | 561 | 590-631 |
| pDRNL:erCER | 458 | 470-512 |
| pDRN:erGFP,<br>pOLEe1:GFP-GUS<br>pPGL1:GFP-GUS<br>E325:GFP<br>pSMR1:nlsGFP-GUS | 488 | 498-535 |
| pDR5:VENUS<br>pCUC2:CUC2:VENUS | 514 | 520-555 |
| SR2200 | 405 | 416-450 |
| DAPI | 405 | 432-455 |

**Extended Data Table 1**
